# The protein and lipid compositions of brain apolipoprotein E particles are cell type-specific

**DOI:** 10.64898/2026.09.07.749850

**Authors:** Evelina Voloviceva, Sandra Goetze, Stephanie Hausler, Safiyo Haji Abukar, Annika Jagels, Alaa Othman, Arnold von Eckardstein, Jérôme Robert

## Abstract

Genetic variation in *APOE*, which encodes a lipid transporter apolipoprotein E (apoE), is the strongest risk factor for late-onset Alzheimer’s disease (AD). Although central nervous system (CNS) apoE is produced by various cell types, including astrocytes, microglia, and pericytes, it is unclear whether apoE lipoproteins are a homogeneous population or if their molecular composition varies depending on their cellular origin. Emerging evidence suggests that CNS apoE lipoproteins contribute to AD-related processes, including amyloid deposition, neuroinflammation, synaptic maintenance, and blood–brain barrier function in a cell type-specific manner.

We hypothesized that the compositions of CNS apoE lipoproteins depend on their cellular origin and investigated the lipid and protein profiles of apoE particles secreted by induced pluripotent stem cell (iPSC)-derived brain cells and derived from human cerebrospinal fluid (CSF) using liquid chromatography-mass spectrometry (LC-MS/MS).

Our results demonstrate that astrocytes, pericytes, and microglia secrete apoE lipoproteins with distinct, cell type-specific lipid and protein profiles. Astrocyte-secreted particles were enriched in sphingomyelins and proteins linked to lipid metabolism, whereas pericyte-secreted particles contained ether-linked phospholipids and proteins associated with antioxidant activity. The lipidation of microglial apoE was markedly lower, but these particles had higher levels of transferrin, apoC-II, and cathepsin D. Notably, one-third of identified features were cell type-specific, highlighting the substantial molecular specialization across cell types. Most lipids and proteins identified in cell-secreted apoE were also detected in CSF-derived particles.

Our findings advance the understanding of apoE biology by shifting the focus from apoE as a single, homogeneous entity to a diverse population of complex particles, the composition of which might be linked to functions in health and neurodegenerative diseases.

## 1. Introduction

Apolipoprotein E (apoE) is the primary structural protein of lipoproteins in the central nervous system (CNS). There, it facilitates lipid transport in cerebrospinal and interstitial fluids and contributes to neuronal maturation, inflammation, and complement regulation, among other processes^1^. Genetic variation in *APOE* is the strongest genetic determinant of sporadic late-onset Alzheimer’s disease (AD) with *APOE4* increasing disease risk, *APOE3* considered neutral, and *APOE2* being protective ^2–4^. While this association is well established, the mechanisms by which apoE contributes to AD remain unclear. A major challenge is that apoE-containing lipoproteins are secreted by multiple CNS cell types and circulate throughout the brain, making it difficult to define cell type-specific functions^5^.

Multiple cell types in CNS, including astrocytes, pericytes, and microglia, produce apoE-containing lipoproteins^5–9^. Knocking out *Apoe* in the whole body reduces amyloid burden and tauopathy in mouse models^10^, however, cell type-specific deletions yield divergent outcomes. Astrocyte-specific *Apoe* knockout decreases amyloid pathology, whereas *Apoe* deletion in microglia does not significantly affect plaque formation, even though microglia are a major source of apoE in the CNS^11,12^. Together, these observations suggest that not all apoE lipoproteins exert the same functions within the CNS. Notably, Yamazaki *et al.* demonstrated that astrocyte-derived apoE lipoproteins are similar in size to pericyte-derived apoE, yet they contain four times less cholesterol^13^. Building on these findings, Merrill *et al.* recently demonstrated that human CSF comprises multiple lipoprotein subspecies composed of hundreds of different proteins with apoE, apoA-I, and clusterin (CLU or apoJ) serving as the major scaffolding proteins. The authors also identified diverse apoE particle-associated proteins involved in the immune response, inflammation, wound healing, and nervous system development^14^.

We therefore hypothesized that the lipid and protein composition of CNS apoE lipoproteins is determined by their cellular origin. Using LC-MS/MS-based lipidomics and proteomics, we characterized apoE particles secreted by human induced pluripotent stem cell (iPSC)-derived astrocytes, microglia, and pericytes. We identified distinct, cell type-specific lipid and protein signatures, which provide new insight into the functional heterogeneity of apoE lipoproteins in the CNS and their potential roles in AD.

## 2. Methods

### 2.1 iPSC culture

BIONi037-A iPSC line (genotyped *APOE3/E3*, derived from a 77-year old healthy female donor) was purchased from The European Bank for induced pluripotent Stem Cells (EBiSC)^15^. iPSCs were cultured on Matrigel-coated (354277 Corning) plates in mTeSR Plus medium (100–0276 StemCell Technologies), passaged every 3-4 days. Culture media changes were performed every 1-2 days. For passaging, cells were washed with PBS (10010023 Gibco), incubated with ReLeSR (100-0483 StemCell Technologies) for 5 min at 37°C with 5% CO_2_, detached in 1 mL culture medium and transferred to new Matrigel-coated plates.

### 2.2 Pericyte differentiation

Pericyte differentiation through the neural crest intermediate stage was carried out as described by Faal *et al*.^16^. Briefly, a single cell suspension of iPSCs was created by treating the cells with Accutase (A1110501 Gibco). Cells were seeded at a density of 20’000 cells per cm^2^ in mTeSR Plus with ROCK inhibitor (ROCKi) (1:1000, 130-103-922 Miltenyi Biotec). The next day, culture medium was changed to the Neural Crest Induction Medium (NCIM): DMEM/F12 (11320-033 Gibco) containing 2% B27 (17504-044 Gibco), 3 µM CHIR 99021 (4423/10 Tocris), 20% BSA (A7906-100G Sigma-Aldrich), 1X Glutamax (35050061 Gibco) and 1:1000 ROCK inhibitor (ROCKi, 130-103-922 Miltenyi Biotec). Medium was changed every day for 6 days (NCIM with 1:1000 ROCKi for the first two days, followed by 4 days with NCIM without ROCKi). On day 6, neural crest cells were passaged with Accutase and seeded at the density of 25’000 cells per cm^2^ in complete Pericyte Medium (1201 ScienCell), containing pericyte growth supplements and 2% FBS following manufacturer’s recommendations. Cells were cultured in Pericyte Medium for the next two weeks. Passages were performed with Accutase and cells were re-plated in media containing 1:1000 ROCKi.

### 2.3 Astrocyte differentiation

Astrocyte differentiation through the neural progenitor intermediate stage was carried out using the STEMdiff SMADi Neural Induction Kit (08581 StemCell Technologies) according to the manufacturer’s instructions. After 3 weeks of neural induction, neural progenitor cells (NPCs) were differentiated to mature astrocytes using the STEMdiff™ Astrocyte Differentiation Kit (100-0013 StemCell Technologies) and STEMdiff™ Astrocyte Maturation Kit (100-0016 StemCell Technologies) following the manufacturer’s instructions.

### 2.4 Microglia differentiation

Microglia were differentiated from iPSCs through the intermediate stage of hematopoietic stem cells using the STEMdiff™ Hematopoietic Kit (05310 StemCell Technologies) and STEMdiff™ Microglia Differentiation Kit (100-0019 StemCell Technologies) according to the manufacturer’s instructions.

### 2.5 Cell marker expression analysis by immunofluorescence staining

Cells were seeded on Matrigel-coated glass coverslips in 24-well cell culture plates (0.08 × 10^6^ cells/well) or in Matrigel-coated black 96-well optical bottom plates (165305 Thermo Scientific) (0.01 × 10^6^ cells/well). After 1-3 days of culture in respective culture media, cells were washed once with PBS and fixed in 4% paraformaldehyde (PFA, 158127 Sigma-Aldrich). After 20 minutes, cells were washed three times with PBS and permeabilized with 0.2% Triton X-100 (X100-1L, Sigma-Aldrich) in PBS. After 10 minutes, cells were washed three times with PBS and blocked in PBS containing 5% Normal Donkey Serum (NDS, S30 Merck) and 0.2% BSA (A4503-50G Sigma-Aldrich) for at least 1 hour. The cells were then incubated overnight at 4°C with the respective primary antibodies, diluted in blocking solution: rabbit anti-Oct4 (RRID: AB_2687916, 1:500), mouse anti-Nanog (RRID: AB_3076592, 1:500), mouse anti-PDGFRβ (RRID: AB_1269704, 1:250), rabbit anti-NG2 (RRID: AB_2877152, 1:200), rabbit anti-S100β (RRID: AB_882426, 1:100), rabbit anti-Iba1 (RRID: AB_2636859, 1:1000), mouse anti-GFAP (RRID: AB_3095092, 1:100), mouse anti-CD45 clone HI30, (RRID: AB_3752289, 1:200).

The next day, cells were washed five times with PBS and incubated at room temperature with the respective donkey anti-mouse AlexaFluor 647 (RRID: AB_162542, 1:500) or anti-rabbit donkey anti-rabbit AlexaFluor 568 (RRID: AB_162542, 1:500) secondary antibodies. After one hour, the cells were washed twice with PBS and cell nuclei were stained with DAPI (MBD0015 Sigma-Aldrich, 1:5000 in PBS) for 7 min before mounting with ProLong™ Gold Antifade Mounting Solution (P36934 Invitrogen). Images were acquired using Cytation 5 cell imaging multimode reader (Agilent BioTek) at 20x or 40x magnification and processed with Gen5 software (Agilent BioTek) and Fiji ImageJ (1.54f, NIH).

### 2.6 Gene expression analysis using RT-qPCR

Total RNA was isolated from respective cells using the NucleoSpin™ Mini Kit for RNA Purification (740955 Macherey Nagel) following manufacturer’s recommendations. 0.5 µg of RNA were reverse-transcribed into cDNA using RevertAid First Strand Synthesis kit (K1621 Thermo Scientific). Real-time qPCR reactions were carried out in a Light Cycler 480-II (Roche) using the LightCycler® 480 SYBR Green I Master Mix (04887352001 Roche) and respective primers for genes of interest: *GAPDH*: for: CCCATGTTCGTCATGGGTGT; rev: TGGTCATGAGTCCTTCCACGATA, *POU5F1*: for: GCAAAGCAGAAACCCTCGTG, rev: GATCTGCTGCAGTGTGGGT, *SOX2*: for: CAACGGCAGCTACAGCATGA, rev: CTCGGACTTGACCACCGAAC, *NANOG*: for: GAGATGCCTCACACGGAGAC, rev: GGGTTGTTTGCCTTTGGGAC, *PAX6*: for: CGCAGGAGGAAGTGTTTTGC, rev: TGCTGATTGGTGATGGCTCA, *NES*: for: TTCCCTCAGCTTTCAGGACCC rev: TGTCTCAAGGGTAGCAGGCA, *SOX1*: for: GGAATGGGAGGACAGGATTT, rev: ACTTTTATTTCTCGGCCCGT, *GFAP*: for: GAGATCGCCACCTACAGGAA, rev: GGCTGGTTTCTCGAATCTGC, *S100B*: for: ACAAGGAAGAGGATGTCTGAGC, rev: GCCGTCTCCATCATTGTCCA, *AQP4*: for: GTGGGGTAAGTGTGGACCTTT, rev: GCCAGAAATTCCGCTGTGAC, *SOX10*: for: ATGAACGCCTTCATGGTGTGGG, rev: CGCTTGTCACTTTCGTTCAGCAG, *PDGFRB*: for: ACAGACTCCAGGTGTCATCC, rev: AGTTGACCACCTCATTCCCG, *MCAM*: for: GCCAGTCCTCATACCAGAGC, rev: TCTTACGAGACGGGGGTAGC, *CSPG4*: for: GTCCTGCCTGTCAATGACCAAC, rev: CTGTGGGTGAGATCATGTGG, *CD34*: for: AACCCAGCCTCCCTCCTAAC, rev: TCCTAGAGAGACGCACCGAG, *SPN*: for: CTGATTCCAGATCCCACGCT, rev: CGGCCTTAGGGTCACTTGTT, *PTPRC*: for: AACAGTGGAGAAAGGACGC, rev: ATGCAGTGGTGTGAGTAGGT, *ITGAM*: for: TGGTGGCTTCCTTGTGGTTC, rev: AAGCCCCTTGCGTTCTCTTG*, TREM2*: for: ACGCTGCGGAATCTACAACC, rev: AGTGGGTGGGAAGGGGATTT.

Gene expression was analyzed with the ΔΔCt method. All results were normalized to GAPDH housekeeping gene. For each marker gene, expression values were normalized (set to 1) to the reference cell type: iPSCs for pluripotency markers; NPC, NC, or Heme for the markers of intermediate stage cells; astrocytes, pericytes, or microglia for target cell markers.

### 2.7 Media collection for the characterization of secreted apoE particles

Terminally differentiated pericytes, astrocytes, and microglia were cultured in their respective delipidated culture media (2% delipidated FBS for pericytes, serum-free media for astrocytes and microglia). After 72 hours with no medium changes, conditioned media were collected and centrifuged at 1000*g* for 10 minutes at 4°C to remove floating cells and debris. Supernatants were transferred to new tubes and stored at 4°C with cOmplete Mini protease inhibitor cocktail tablets (11 836 153 001 Roche, 1 tablet per 10 mL media) for up to one week before apoE concentration determination by ELISA and apoE particle immunoprecipitation.

### 2.8 CSF preparation for apoE immunoprecipitation

Three anonymized samples of human CSF were purchased from BioIVT (product number HUMANCSFR-0101364) and pooled together. The pooled sample was centrifuged at 1000*g* for 10 minutes at 4°C to remove debris. The supernatant was transferred to a new tube and used immediately for apoE concentration determination by ELISA and apoE particle immunoprecipitation.

### 2.9 ApoE concentration determination using ELISA

The concentration of apoE in CSF and cell-conditioned media was determined by apoE ELISA as previously described^17^. Briefly, high binding 96-well plates (3855 Thermo Scientific) were coated with 1.67 µg/mL apoE monoclonal antibody E276 (3712-3 Mabtech) in PBS at 4°C. After overnight incubation, plates were washed with PBST (PBS + 0.05% Tween-20 (P3563 Sigma-Aldrich) and blocked with 0.1% Blocker A (R93BA-4 Meso Scale Discovery) diluted in PBST. 100 µL of samples or human recombinant apoE (3712-10 Mabtech) standards (0.49 - 500 ng/mL) diluted in PBST were added to the wells and incubated at room temperature. After one hour, biotinylated anti-human apoE monoclonal E887 antibody (3712-6-250 Mabtech) and Streptavidin-HRP (3310-9 Mabtech) were used for detection. Peroxidase activity was detected with QuantaBlu Fluorogenic Peroxidase Substrate Kit (15169 Thermo Scientific) according to manufacturer’s instructions. Fluorescence readings (Ex 325 nm/Em 420 nm) were performed with Cytation 5 microplate reader (Agilent BioTek). Sigmoidal, 4-parameter logistic curve fitting was performed in GraphPad Prism to determine apoE concentrations.

### 2.10 Immunoprecipitation of apoE particles

Antibody-bead conjugates were prepared with the Dynabeads® Antibody Coupling Kit (14311D Thermo Scientific) according to the manufacturer’s instructions. Goat anti-human apoE antibody (A81-118A-1MG Fortis) and Dynabeads® M-270 Epoxy (14301 Thermo Scientific) were used per one coupling reaction at a ratio of 35 µg to 5 mg, respectively. At the end of the protocol, antibody-bead conjugates were resuspended in the storage buffer (SB) and stored at 4°C for up to four days. On the day of the immunoprecipitation, antibody-bead conjugates were washed six times with PBS (10010023 Gibco) to remove traces of detergents from the storage buffer. Before apoE immunoprecipitation, the cell-conditioned media were concentrated using Amicon Ultra 30 kDa centrifugal filters (UFC5030 Merck Millipore) following the manufacturer’s instructions. Astrocyte-conditioned media were added to the antibody-bead conjugates with volumes adjusted to contain 150 ng apoE per reaction. Other cell-conditioned media and CSF were added to the antibody-bead conjugates with volumes adjusted to contain 500 ng apoE per one reaction. Equal final reaction volumes were maintained across all samples by adjusting sample input volumes with PBS (for CSF) or vehicle medium (for conditioned media). Samples were incubated overnight (20 hours) at 4°C with end-over-end rotation. After 20 hours, supernatants were removed using a magnetic separation rack. Antibody-bead conjugates with immunoprecipitated apoE were washed three times with cold PBS (15 min end-over-end rotation with each wash). Immunoprecipitates on beads were stored at -70°C until further processing.

### 2.11 Proteotype determination

The analysis included immunoprecipitated apoE from 1) human CSF, 2) astrocyte-conditioned media, 3) pericyte-conditioned media, 4) microglia-conditioned media, and 5) iPSC-conditioned media. A mix of all cell culture media (vehicle media without cell contact, in equal volumes) was used as the negative control. The control samples were processed in the same manner as the experimental samples, including incubation with the antibody-bead conjugates.

Immunoprecipitated apoE particles on magnetic beads were lysed and processed with iST 8x kits (P.O.00001 PreOmics) to obtain the peptide mix prior to analysis by liquid chromatography coupled to mass spectrometry (LC-MS/MS).

An estimated 500 ng of peptides, as quantified by NanoDrop, were injected per sample for LC-MS/MS analysis. Peptides were separated on a Vanquish™ Neo UHPLC system (Thermo Scientific) equipped with a 50 cm μPAC™ Neo HPLC column (COL-nano050NeoB, Thermo Scientific). Samples were loaded at 0.7 μL min⁻¹ in 100% mobile phase A (99.9% H₂O, 0.1% formic acid) with a maximum pressure of 400 bar. Peptides were eluted at 300 nL min⁻¹ using a stepped gradient from 8–32% mobile phase B (80% acetonitrile, 0.1% formic acid) over 36 min, followed by 32–50% B over 4 min. Samples were analyzed in technical triplicates by data-dependent acquisition (DDA) on an Orbitrap Exploris™ 480 mass spectrometer (Thermo Scientific). Full MS scans were acquired over an *m/z* range of 350–1200 at a resolution of 120,000, with a normalized AGC target of 300% and a maximum injection time of 25 ms. The top 20 precursor ions were fragmented by HCD (normalized collision energy, 30%), and MS/MS spectra were acquired at a resolution of 15,000, using a normalized AGC target of 50%, maximum injection time set to auto, and a dynamic exclusion of 15 s.

Raw files were analyzed with FragPipe v23.0 using the DDA+ workflow. Spectra were searched with MSFragger v4.3 against the reviewed UniProt Homo sapiens reference proteome (June 2025) supplemented with common contaminants and reverse decoy sequences. The search was performed with strict trypsin specificity, allowing for up to two missed cleavages and a precursor and fragment mass tolerance of 20 ppm. Carbamidomethylation of cysteine was set as a fixed modification, while methionine oxidation and protein N-terminal acetylation were included as variable modifications. Peptide-spectrum matches were rescored using MSBooster with DIA-NN-predicted spectra and retention times, followed by validation with Percolator. Protein inference was performed with ProteinProphet, and peptide and protein identifications were filtered to a 1% false discovery rate (FDR). Label-free quantification was performed with IonQuant using the MaxLFQ algorithm with match-between-runs enabled.

The initial protein list was curated by removing likely contaminants^18^: keratins, histone, ribosome, heat shock protein subunits, and fragments of the antibody used for the immunoprecipitation. Extracellular matrix, cytoskeletal proteins, and enzymes were retained despite their classification as frequent background proteins in proteomics studies^18^ because of their possible biological relevance and prior detection in CSF lipoproteins by Merrill *et. al.* in 2023^14^. Only proteins enriched (fold change (FC) ≥ 2 and adjusted *P* value < 0.05) in each experimental condition compared to vehicle media controls were defined as associated with apoE particles. Protein filtering criteria and results are included in the Supplementary File 1. STRING database was used to assign proteins to GO terms and identify potential protein functions^19^.

### 2.12 Lipidome determination

For lipidomics analyses, we included the same conditions as for proteomics.

Lipid extraction from the immunoprecipitated apoE particles on magnetic beads was carried out using the butanol:methanol (BuMe, 1:1, v/v) method prior to LC-MS/MS analysis. 200 µL of BuME, including the internal standard mixture EquiSPLASH (Avanti Polar Lipids, 1 µg/sample), were added per sample. The samples were vortexed and briefly sonicated before shaking for 20 min in a thermomixer at 1000 rpm at room temperature (RT). All the subsequent steps of the extraction were performed at RT. Samples were centrifuged for 10 min at 16’000 rcf, the supernatants were transferred to new tubes and dried under a gentle nitrogen stream. Then, the samples were reconstituted in 100 µL of MeOH:IPA (1:1, v/v), vortexed and shaken in a thermomixer for 10 min, 1000 rpm. Lastly, the samples were centrifuged for 10 min at 16’000 rcf and the supernatants were transferred to LC-MS glass vials.

Lipid profiling analysis was performed as previously described^20^ with modifications. Analyses were conducted on a Thermo Vanquish Horizon Binary Pump UHPLC system coupled to a Thermo Orbitrap Exploris 240 mass spectrometer with a heated electrospray ionization (HESI) source. Chromatographic separation was achieved on a Waters ACQUITY Premier BEH C18 column (1.7 µm, 2.1 × 50 mm) maintained at 60 °C. Mobile phase A was MeCN:H_2_O (60:40, v/v) with 5 mM ammonium acetate, and mobile phase B was MeCN:IPA (10:90, v/v) with 5 mM ammonium acetate. The following gradient was applied at 0.6 mL min⁻¹: 0 min, 15% B; 1.87 min, 30% B; 2.08 min, 48% B; 9.17 min, 82% B; 9.59 min, 99% B; 10.08– 12.6 min, 99% B; followed by re-equilibration to 15% B. The injection volume was 3 µL. The Orbitrap Exploris 240 was operated in positive ionization mode with the following source settings: sheath gas = 30, auxiliary gas = 13, spray voltage = 3,800 V, vaporizer temperature = 350 °C, ion transfer tube temperature = 300 °C. Internal mass calibration was performed at the start of each run, and mild trapping was enabled to minimize in-source fragmentation. MS1 settings: resolution 60,000; scan range 150–2000 m/z; AGC target 1e6; maximum injection time 10 ms. MS2 settings: DDA of top five precursors, isolation window 1 m/z; stepped normalized collision energy 10, 30, and 40%; resolution 30,000; AGC target 1e5; maximum injection time 10 ms; RF lens 50%. Pooled quality controls (QC) of all samples were used in the beginning, middle, and end of the batch.

Data analysis and lipid identification were performed using the Compound Discoverer software (Thermo Scientific, version 3.3.3.200). Lipid identification was achieved by comparing all lipids with a lipid blast *in silico* library result and confirming the match by the presence of diagnostic fragments according to the Lipidomics Standards Initiative (LSI) guidelines^21^. Only lipids that matched the expected fragmentation spectra, mass accuracy threshold, and the expected pattern of retention times were included in the analyses. Only lipids enriched (fold change (FC) ≥ 2 and adjusted *P* value < 0.05) in each experimental condition compared to vehicle media controls were defined as associated with apoE particles. Lipid filtering criteria and results are included in the Supplementary File 2.

### 2.13 Statistics

Normality or lognormality of data distribution were assessed with the Shapiro-Wilk test. Comparisons between two groups were performed using Welch’s t-test, comparisons between multiple groups — with one-way ANOVA followed by Dunnet’s post hoc multiple comparisons test. Mann-Whitney and Kruskal-Wallis tests were used as non-parametric alternatives for two-group and multiple group comparisons, respectively. Data visualization and statistical analyses were conducted in GraphPad Prism and R software. ChatGPT (model GPT-4o, OpenAI) and Claude (model Sonnet 5, Anthropic) Large Language Models were used for R code troubleshooting and optimization.

Lipidomics and proteomics data analysis was carried out in R using Bioconductor packages *lipidr* and *limma*. Data were visualized with *pheatmap*, *ggplot2,* and *eulerr* packages. Lipidomics data were normalized using internal standards and log_2_-transformed. Relative lipid abundances were calculated from internal standard-normalized linear data, before log_2_-transformation. Absolute concentrations were determined by normalizing analyte signal to internal standards of known concentration.

Proteomics data were log_2_-transformed and median-centered. Relative protein abundances were calculated from median-centered linear data, before log_2_-transformation. Proteins with at least 40% non-missing values across all samples were retained for analysis. Missing values were imputed using the minimum probability (MinProb) method from the *imputeLCMD* R package. For heatmap and volcano plot visualization, proteomics data were additionally normalized using APOE intensity after median-centering. *P* values were adjusted for multiple testing with the Benjamini-Hochberg method to control the false discovery rate (FDR). Differences were considered statistically significant at p < 0.05, FDR < 0.05, and |FC| ≥ 2 (|log2FC| ≥ 1), where applicable.

For all analyses, one replicate represented one well derived from an independent differentiation. Three replicates were analyzed per cell type for -omics analyses and at least three replicates per cell type for qPCR.

## 3. Results

### 3.1 *APOE3/E3* iPSCs differentiate into apoE-secreting brain cells

To investigate the composition of apoE particles secreted by different brain cell types, we differentiated human iPSCs into astrocytes, pericytes and microglia (Fig. 1 A). First, we verified the pluripotency of iPSCs by positive expression of pluripotency markers. Gene expression of *POU5F1* (encoding Oct-4)*, SOX2, NANOG* was confirmed by RT-qPCR (Sup. Fig. 1 A, C, E) and Oct-4 and Nanog protein expression – by immunofluorescence (IF) staining (Fig.1 B). We confirmed the differentiation of iPSCs into target cell types by analyzing the expression of cell type-specific markers on the RNA and protein levels. We included iPSCs, intermediate-stage cells, and terminally differentiated cells in the gene expression analyses.

**Figure 1.**
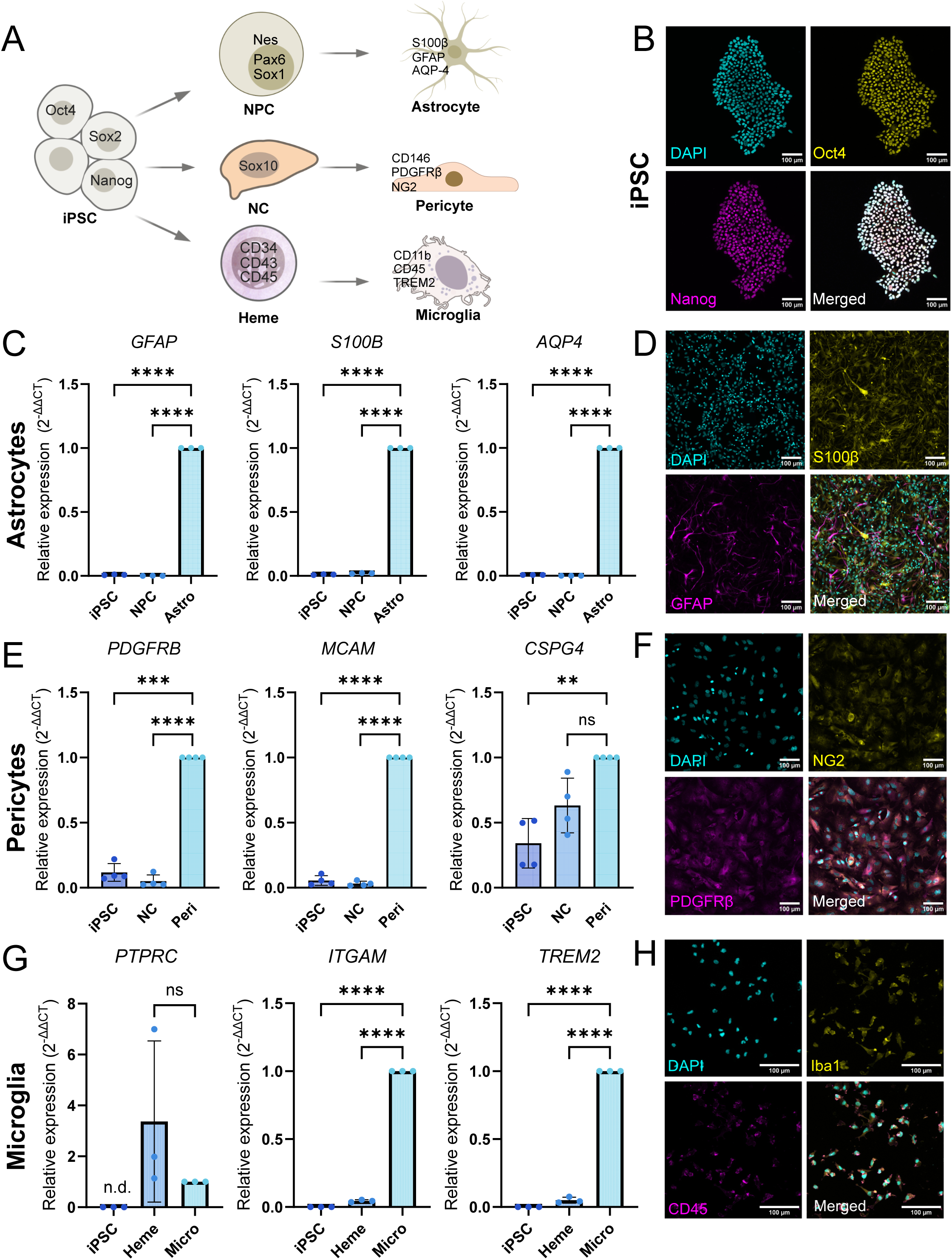
Differentiation of iPSC into pericytes, astrocytes and microglia. A) Schematic differentiation of iPSCs to astrocytes, pericytes, and microglia. Astrocytes (markers: S100β, GFAP, AQP-4) were differentiated via Neural Progenitor cells (NPC, markers: Pax6, Sox1, Nestin (Nes)); Pericytes (markers: CD146, PDGFRβ, NG2) were differentiated via Neural Crest cells (NC, marker Sox10); Microglia (markers: CD11b, TREM2, CD45^low^) were differentiated via Hematopoietic stem cells (Heme, markers: CD34, CD43, CD45). B) Expression of pluripotency markers (Oct4, Nanog) in iPSCs, nuclei counterstained with DAPI. Scale bars 100 µm. C) Gene expression of *GFAP, S100B, AQP4* in iPSCs, NPC, and astrocytes by qRT-PCR. D) Expression of S100β and GFAP in astrocytes. Nuclei counterstained with DAPI. Scale bars 100 µm. E) Gene expression of pericyte markers (*PDGFRB, MCAM* – encoding CD146*, CSPG4* – encoding NG2) in iPSCs, NC, and pericytes by qRT-PCR. F) Expression of NG2 and PDGFRβ in pericytes. Nuclei counterstained with DAPI. Scale bars 100 µm. G) Gene expression of microglial markers (*PTPRC* – encoding CD45*, ITGAM* – encoding CD11b*, TREM2*) in iPSCs, Heme, and microglia by qRT-PCR. H) Expression of Iba1 and CD45 in microglia. Nuclei counterstained with DAPI. Scale bars 100 µm. Points in graphs represent individual experiments (biological replicates, n = 3-4), bars represent the mean, and error bars ± SD, *p < 0.05, **p < 0.01, ***p < 0.001, ****p < 0.0001, ns – not significant, n.d.-not detected within 40 amplification cycles. iPSC – induced pluripotent stem cells, NC – neural crest, Heme – hematopoietic stem cells, Peri – pericytes, Micro – microglia, Astro – astrocytes.

Astrocytes were differentiated via neural progenitor cells (NPC). Expression of astrocyte markers *GFAP*, *S100B*, and *AQP4* (Fig. 1 C) together with loss of pluripotency markers *POU5F1* and *NANOG* (Sup. Fig. 1 A) in the resulting cells confirmed differentiation into astrocytes. iPSC-derived astrocytes conserved the expression of *SOX2* and the NPC markers *PAX1*, *NES*, and *SOX1* (Sup. Fig. 1 A-B), suggesting that a population of cells that are incompletely differentiated remained in culture at the end of the differentiation protocol. We further confirmed the expression of S100β and GFAP proteins in iPSC-derived astrocytes (Fig. 1 D).

Pericytes were differentiated via neural crest cells (NC). Expression of pericyte markers *PDGFRB*, *MCAM* (encoding CD146), and *CSPG4* (encoding NG2) (Fig. 1 E), together with loss of pluripotency markers *POU5F1*, *SOX2*, and *NANOG* (Sup. Fig. 1 C) and loss of NC marker *SOX10* (Sup. Fig. 1 D) in differentiated cells confirmed differentiation into pericytes. iPSC-derived pericytes also expressed NG2 and PDGFRβ on the protein level (Fig. 1 F).

Microglia were differentiated via hematopoietic stem cells (Heme). Expression of microglial markers *ITGAM* (encoding CD11b) and *TREM2* (Fig. 1 G), together with loss of pluripotency markers *POU5F1*, *SOX2* and *NANOG* (Sup. Fig. 1 E) and downregulation of hematopoietic stem cell markers *CD34* and *SPN* (encoding CD43) (Sup. Fig. 1 F) in the resulting cells confirmed differentiation into microglia. *PTPRC* (encoding CD45) expression was not detected in iPSCs, but it was present in resulting iPSC-derived microglia. However, *PTPRC* expression was not significantly higher in differentiated microglia than in hematopoietic stem cells, suggesting that the resulting cells likely belong to the CD45^low^, CD11b^+^ population of brain microglia (Fig. 1 G)^22^. iPSC-derived microglia also expressed the Iba1 and CD45 proteins (Fig. 1 H). Finally, we confirmed that iPSC-derived astrocytes, pericytes, and microglia, as well as undifferentiated iPSCs, expressed and secreted apoE (Sup. Fig. 2 A).

**Figure 2.**
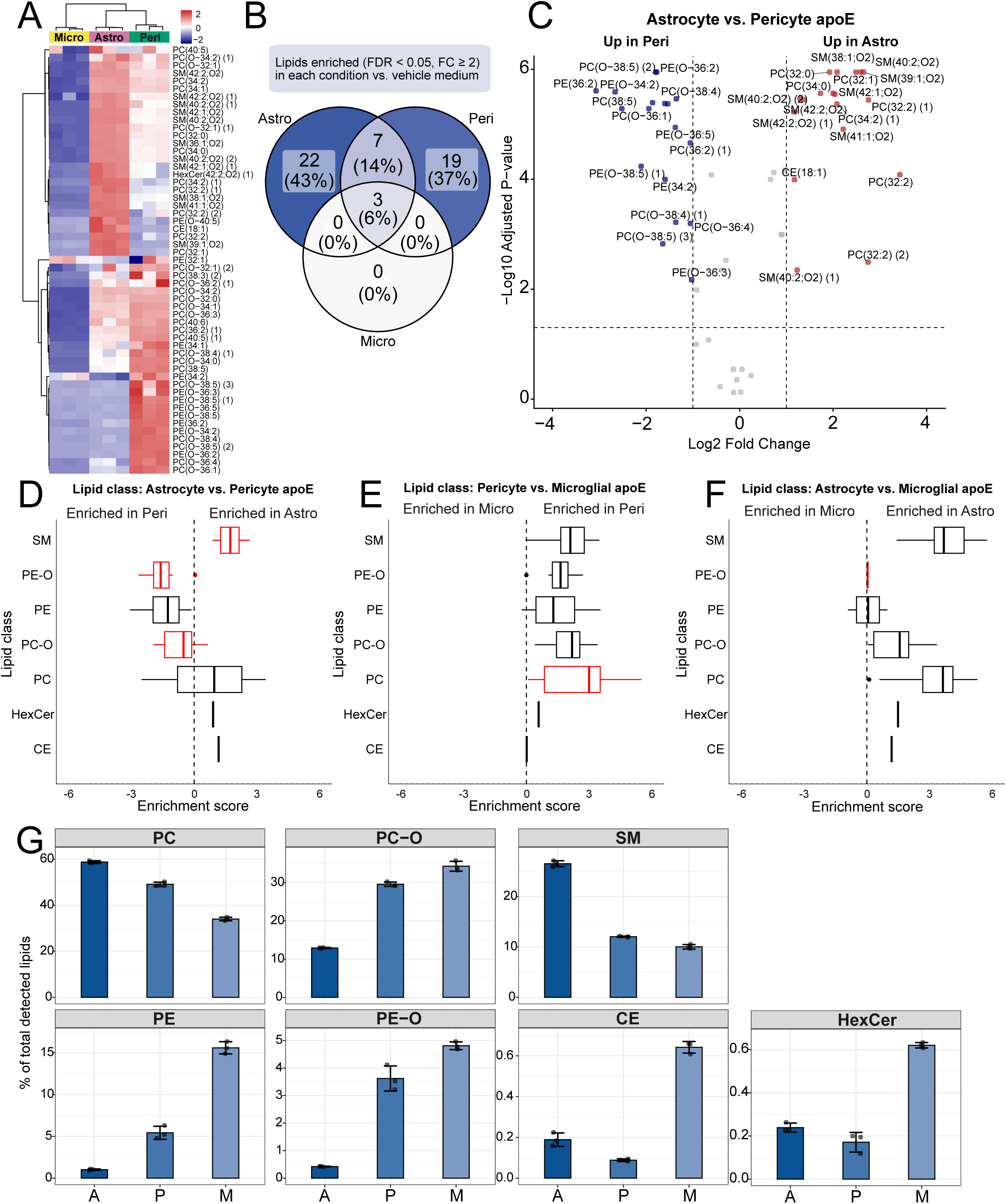
Lipid profiling reveals that different lipids are associated with apoE particles depending on the cell type of origin. A) Heatmap represents color-coded abundance (Z-scored, row-scaled) of lipid species associated with apoE particles secreted by microglia, astrocytes, and pericytes. Only lipids enriched (log_2_FC ≥ 1, FDR < 0.05) in each condition vs. vehicle media controls are shown. B) The overlap of lipids identified as associated with astrocyte, pericyte, and microglial apoE, defined as enriched (log_2_FC ≥ 1, FDR < 0.05) in each condition vs. vehicle media controls. C) Differential expression of lipids associated with astrocyte vs. pericyte apoE particles. Each point represents one lipid species. Significance thresholds: |log_2_FC| ≥ 1, FDR < 0.05. D-F) Lipid class enrichment analysis of differentially abundant lipids in astrocyte vs. pericyte (D), pericyte vs. microglial (E), and astrocyte vs. microglial (F) in apoE particles. Lipid class distributions are shown as boxplots with medians as center lines, whiskers extend to the extreme non-outlier values, dots represent outliers. Significant classes (FDR < 0.05) are highlighted. FC - fold change, FDR – false discovery rate. G) Relative abundance of lipid classes in astrocyte-, pericyte-, and microglia-secreted apoE particles. Points in graphs represent individual replicates (n = 3), bars represent the mean, and error bars ± SD. Astro or A – astrocyte-, Peri or P – pericyte-, Micro or M – microglia-secreted apoE.

### 3.2 Lipids associated with brain apoE differ depending on the cell type of origin

To characterize the lipid composition of apoE lipoproteins secreted by iPSC-derived astrocytes, pericytes, and microglia, we maintained the cells in respective delipidated culture media for three days before immunoprecipitation and lipidomics analysis of the secreted apoE particles. We profiled sphingolipids, phospholipids, glycerolipids, and cholesteryl esters.

The comparison of astrocyte-, pericyte-, and microglia-secreted apoE particles revealed different and unique lipidation patterns depending on the cell type of origin. Microglia-secreted apoE were poorly lipidated, while astrocyte- and pericyte-secreted particles were enriched in several lipid classes (Fig. 2 A).

We then performed differential lipid expression analysis, comparing apoE from each brain cell type to vehicle media controls, and defined cell type-specific apoE lipids as those significantly enriched in that comparison. 22 of 51 lipid species present in brain cell-secreted apoE (43%) were unique to astrocyte-secreted particles (8 phosphatidylcholines (PC), 11 sphingomyelins (SM), hexosylceramide HexCer 42:2;O2, cholesteryl ester CE 18:1, and ether-linked phosphatidylcholine PC O-32:1). 19 lipid species (37%) were unique to pericyte-secreted apoE (5 PC, 6 PC-O, 2 phosphatidylethanolamines (PE), and 6 ether-linked PE (PE-O)). Seven lipid species (PC O-34:1, PC O-36:3, PC O-34:0, PC O-32:0, PC 32:2, PC O-38:4, PC O-36:2)) were shared among astrocyte- and pericyte-secreted particles. Three lipid species were detected in apoE particles of all three brain cell types, namely PC O-34:2 and the two isobars of PC O-32:1. None were unique to microglia-secreted apoE (Fig. 2 B, Sup. Table 1).

Differential expression analysis confirmed that astrocyte-secreted apoE particles are enriched in SM, while pericyte-secreted apoE particles are enriched in ether-linked phosphoglycerides: PC-O and PE-O (Fig. 2 C). The most differentially abundant lipid species were SM 38:1;O2, SM 40:2;O2, and SM 39:1;O2 in astrocyte-secreted apoE and PC O-38:5, PE O-36:2, and PE O-34:2 in pericyte-secreted particles. Lipid class enrichment comparison between astrocyte- and pericyte-secreted apoE particles further confirmed the enrichment of SM or PC-O and PE-O, respectively (Fig. 2 D). Lipid class enrichment analyses of pericyte- vs. microglia-secreted particles (Fig. 2 E) and astrocyte- vs. microglia-secreted apoE particles (Fig. 2 F) showed enrichment of almost all lipid classes in either astrocyte or pericyte apoE particles, confirming poor lipidation of microglial apoE.

We further investigated the composition of apoE particles secreted by each brain cell type using relative abundance calculations for each lipid class. PC 40:5 and its one isobaric form were removed from relative abundance calculations as likely cell culture artifacts due to their high abundance in cell-secreted apoE – 10.7% of total lipid signal in astrocyte-, 20.6% in pericyte-, and 54.4% in microglia-secreted particles, but not CSF apoE, where it comprised 0.6% of total lipid signal (Sup. File 3). PC, PC-O, and SM accounted for the largest proportions of total detected lipids across investigated cell types. However, the abundance of each lipid class differed between pericyte-, astrocyte-, and microglia-secreted particles. Notably, pericyte-secreted apoE particles contained more PC-O than astrocyte-secreted particles (29.6% vs. 12.9% respectively) and less SM (12.0% vs. 26.5% respectively). The lipid class composition of microglia-secreted apoE resembled pericyte-secreted apoE particles in PC-O and SM levels but microglial apoE contained more PE (15.6% vs. 5.4% in pericyte- and 1.0% in astrocyte-secreted particles) and PE-O (4.8% vs. 3.6% in pericyte- and 0.4% in astrocyte-secreted apoE particles) (Fig. 2 G). CE and HexCer accounted for less than 1% of total detected lipids across the investigated cell types. These lipid classes were relatively most abundant in microglia-secreted particles, but their absolute concentrations were highest in astrocyte-secreted apoE particles and similar between pericytes and microglia (Fig. 2 G, Sup. Fig. 3 A). Overall, we demonstrated that the lipid composition of apoE lipoproteins depends on their cellular origin. Particles derived from astrocytes, pericytes, and microglia showed different lipidation patterns and cell type-specific lipid enrichment.

**Figure 3.**
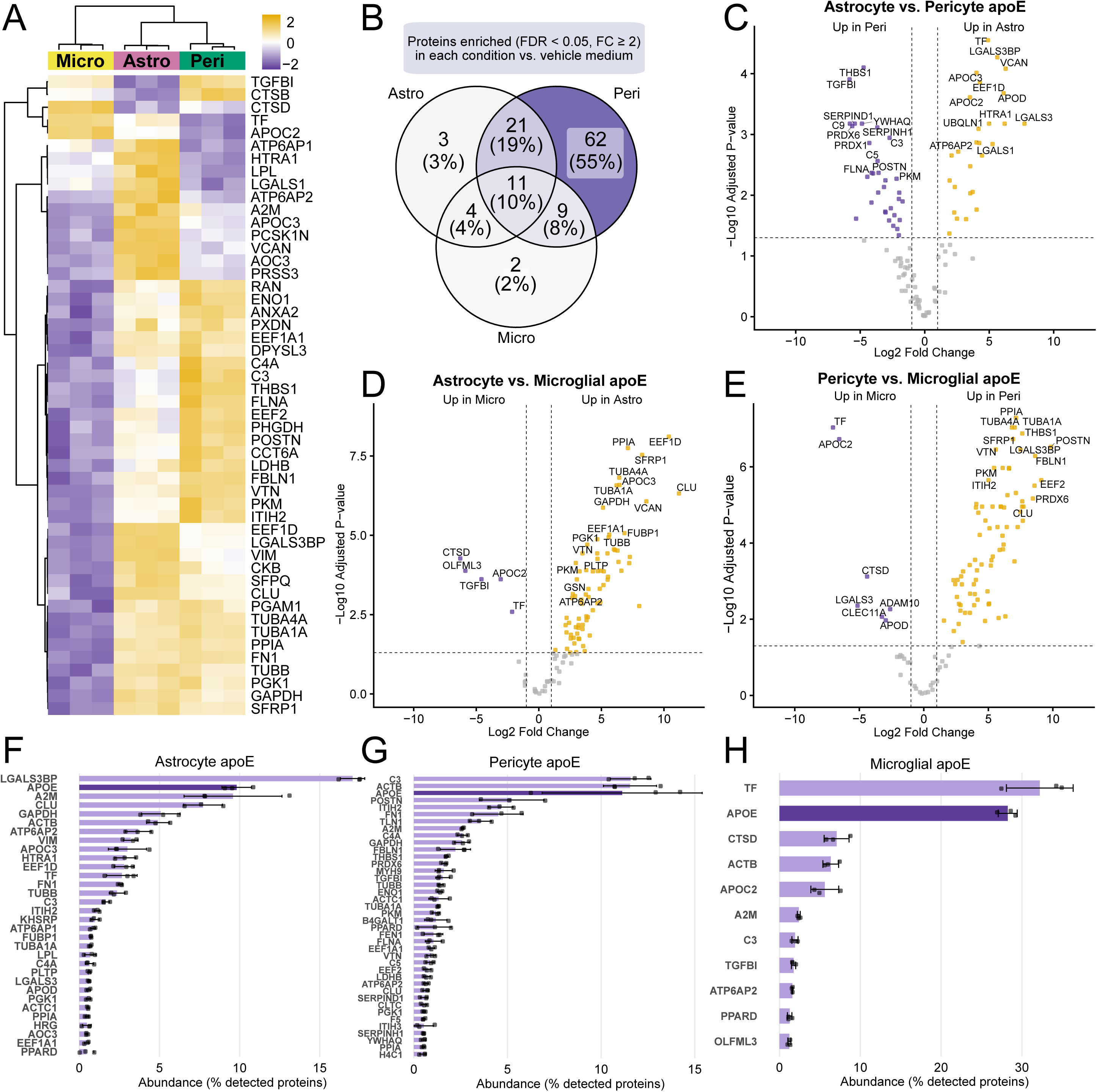
Proteotype profiling reveals that different proteins are associated with apoE particles depending on the cell type of origin. A) Heatmap represents color-coded abundance (Z-scored, row-scaled) of proteins associated with apoE particles secreted by microglia, astrocytes, and pericytes. Top 50 proteins ranked by ANOVA F-statistic are shown. Only proteins enriched (log_2_FC ≥ 1, FDR < 0.05) in each condition vs. vehicle media controls were included in the analysis. Data are normalized to APOE intensity in addition to global (median-centered) normalization. B) The overlap of proteins identified as associated with astrocyte, pericyte, and microglial apoE, defined as enriched (log_2_FC ≥ 1, FDR < 0.05) in each condition vs. vehicle media controls. C-E) Differential expression of proteins associated with astrocyte vs. pericyte (C), astrocyte vs. microglial (D), and pericyte vs. microglial (E) apoE particles. Data are normalized to APOE intensity in addition to global (median-centered) normalization. Significance thresholds: |log_2_FC| ≥ 1, FDR < 0.05. Proteins contributing cumulatively to 90% of total astrocyte-(F), total pericyte- (G), and total microglia-secreted (H) apoE proteotype and their relative abundances, expressed as percentage of all detected proteins. Points in graphs represent individual replicates (n = 3), bars represent the mean, and error bars ± SD. Astro – astrocyte-, Peri – pericyte-, Micro – microglia-secreted apoE.

### 3.3 Proteins associated with brain apoE differ depending on the cell type of origin

To investigate the protein compositions of brain cell-secreted apoE lipoproteins, we cultured iPSC-derived cells as described above and immunoprecipitated secreted apoE particles for proteotype analysis. The comparison of microglia-, astrocyte-, and pericyte-secreted apoE particles revealed different and unique protein expression patterns depending on the cell type of origin (Fig. 3 A). Among detected features, we observed several proteins linked to lipid metabolism (e.g., phospholipid transfer protein – PLTP, clusterin – CLU, apolipoprotein C-II – APOC2)^23^, extracellular matrix (ECM) components or ECM-associated proteins (e.g., nidogen-1 – NID1, transforming growth factor-beta-induced protein – TGFBI, annexin A2 – ANXA2)^24^, proteins involved in amyloid processing (CLU, cathepsin D – CTSD, disintegrin and metalloproteinase domain-containing protein 10 – ADAM10)^25–27^ or amyloid formation (lactadherin – MFGE8)^28,29^, and ATPase-associated proteins, including ATP6AP2 – a subunit of the vacuolar ATPase responsible for lysosomal acidification^30^. (Sup. Table 2). Similarly to lipids, many proteins enriched in astrocyte- and pericyte-secreted particles were downregulated in microglia-secreted apoE (Fig. 3 A).

We then defined cell type-specific apoE proteins via differential expression analysis, comparing each cell type’s apoE to vehicle media controls. The overlap of proteins revealed that 11 of 112 (10%) brain cell-secreted proteins associated with apoE were shared between astrocyte-, pericyte, and microglia-secreted particles. These included APOE, ATP6AP2 and ATP6AP1, galectin-1 (LGALS1), ADAM10, osteoponin (SPP1), serine protease HTRA1, NID1, profilin-1 (PFN1), serine/threonine-protein phosphatase PPP2R1A, and ubiquilin-1 (UBQLN1). STRING GO database search suggested a functional link between APOE and ADAM10, SPP1, LGALS1, and UBQLN1^19^. 62 (55%) of identified proteins were uniquely associated with pericyte-secreted apoE particles. A STRING GO database search revealed that these proteins are linked to antioxidant and peroxidase activities, as well as cell adhesion molecule binding^19^. Additionally, we observed complement system components C3, C4A, C5, and C9 among pericyte-associated apoE particle proteins. Twenty-one (19%) of all brain cell-secreted proteins associated with apoE were found in both pericyte- and astrocyte-secreted apoE particles, including ECM components (e.g., fibronectin-1 – FN1), glycolytic enzymes (e.g., glyceraldehyde-3-phosphate dehydrogenase – GAPDH and pyruvate kinase – PKM), and lipid metabolism regulators (e.g., PLTP and CLU). We noted three proteins (versican – VCAN, proprotein convertase subtilisin/kexin type 1 inhibitor – PCSK1N, and apolipoprotein D – APOD) uniquely associated with astrocyte-secreted particles and four proteins shared between astrocyte and microglial apoE (lipoprotein lipase – LPL, osteolectin – CLEC11A, galectin-3 – LGALS3, and APOC2). Nine proteins were detected in both pericyte- and microglia-secreted particles (ATPase subunit ATP5F1B, beta-1,4-galactosyltransferase 1 – B4GALT1, ganglioside activator GM2A, tensin-4 – TNS4, olfactomedin-like protein 3 – OLFML3, peroxisome proliferator-activated receptor delta – PPARD, GTP-binding nuclear protein RAN, TGFBI, CTSD) and two proteins were uniquely associated with microglial apoE (transferrin – TF, and coiled-coil domain-containing protein CCDC40) (Fig. 3 B, Sup. Table 2).

Differential expression analysis of astrocyte- vs. pericyte-secreted particles revealed the enrichment of TF, LGALS3 and its binding protein (LGALS3BP), VCAN, APOC2, and APOC3 in astrocyte-secreted apoE particles, while the expression of proteins with peroxidase activity (peroxiredoxins 6 and 1 – PRDX6 and PRDX1)^31^, complement components, and ECM-binding proteins TGFBI and thrombospondin-1 (THBS1) was higher in pericyte-secreted apoE particles (Fig. 3 C).

Differential expression analyses of astrocyte- vs. microglia-secreted particles (Fig. 3 D) and pericyte- vs. microglia-secreted apoE particles (Fig. 3 E) confirmed that microglia-secreted apoE particles contained less proteins in comparison to astrocyte- and pericyte-secreted particles. Peptidylprolyl isomerase A (PPIA), also known as cyclophilin A (CypA), was highly enriched in both astrocyte- and pericyte-secreted apoE compared to microglia-secreted apoE. Gelsolin (GSN) was enriched in astrocyte-secreted particles. Proteins associated with the extracellular matrix (e.g., THBS1, fibulin-1 – FBLN1, periostin – POSTN, vitronectin – VTN)^24^ and PRDX6 were more represented in pericyte apoE (Fig. 3 E). However, TF, proteins potentially involved in amyloid processing (CTSD, TGFBI, ADAM10)^19^, and APOC2 were enriched in microglia-secreted particles (Fig. 3 D-E).

Relative abundance calculations of proteins associated with astrocyte-secreted apoE particles revealed that 18 proteins contribute cumulatively to 90% of their proteotype. LGALS3BP was the most abundant (17.0%), followed by APOE (9.8%), alpha-2-macroglobulin (A2M; 9.6%) and CLU (7.7 %) (Fig. 3 F). In pericyte-secreted apoE particles, 20 proteins contributed to 90% of the proteotype, with C3 (11.6%), beta-actin (ACTB; 11.5%), and APOE (11.1%) as the most abundant (Fig. 3 G). In microglia-secreted apoE particles, only 11 proteins contributed to the 90% of the proteotype, with TF (32.2%) and APOE (28.3%) being the most abundant, followed by CTSD (7.1 %) (Fig. 3 H, Sup. File 4). Together, these results demonstrate that each brain cell type secretes apoE particles with a unique protein composition.

### 3.4 iPSC secrete apoE particles with distinct lipid and protein compositions

Although the primary focus of this study was the composition of brain cell-derived apoE lipoproteins, we also observed apoE secretion by undifferentiated human iPSCs (Sup. Fig. 2 A). We therefore investigated the lipidome and proteotype of apoE particles secreted by iPSCs. These particles were characterized by high proportion of PE (56.3%), followed by PC-O (19.4%), and PC (18.3%). SM were the least abundant in iPSC-secreted apoE particles compared to apoE secreted by other cell types (2.5 % of total lipid signal) (Sup. Fig. 4 A, Sup. Fig. 3 B, Sup. File 3). APOC3 was highly represented in iPSC-secreted particles (39.0% of total protein signal), followed by APOE (22.5%) and GAPDH (5.4%) (Sup. Fig. 4 E, Sup. File 4). Differential expression analysis of iPSC-secreted and astrocyte-, pericyte- or microglia-secreted apoE particles confirmed the enrichment of PE and ether-linked PE (PE-O), as well as the depletion of SM and PC in iPSC-secreted apoE particles (Sup. Fig. 4 B-D). Although we observed similar protein groups (proteins linked to lipid metabolism, extracellular matrix, peroxidase activity, amyloid beta processing) associated with iPSC-secreted apoE particles, differential expression analysis identified distinct protein abundance profiles in iPSC apoE compared to apoE secreted by different brain cell types. APOC3 was one of the most highly upregulated proteins in iPSC-secreted apoE in comparison to astrocyte-, pericyte-, and microglia-secreted particles. Additionally, peroxiredoxins, ECM proteins OLFML3 and VCAN, C9, and MFGE8 were among the most upregulated features in iPSC-secreted particles compared to brain cell-secreted apoE (Sup. Fig. 4 F-H).

**Figure 4.**
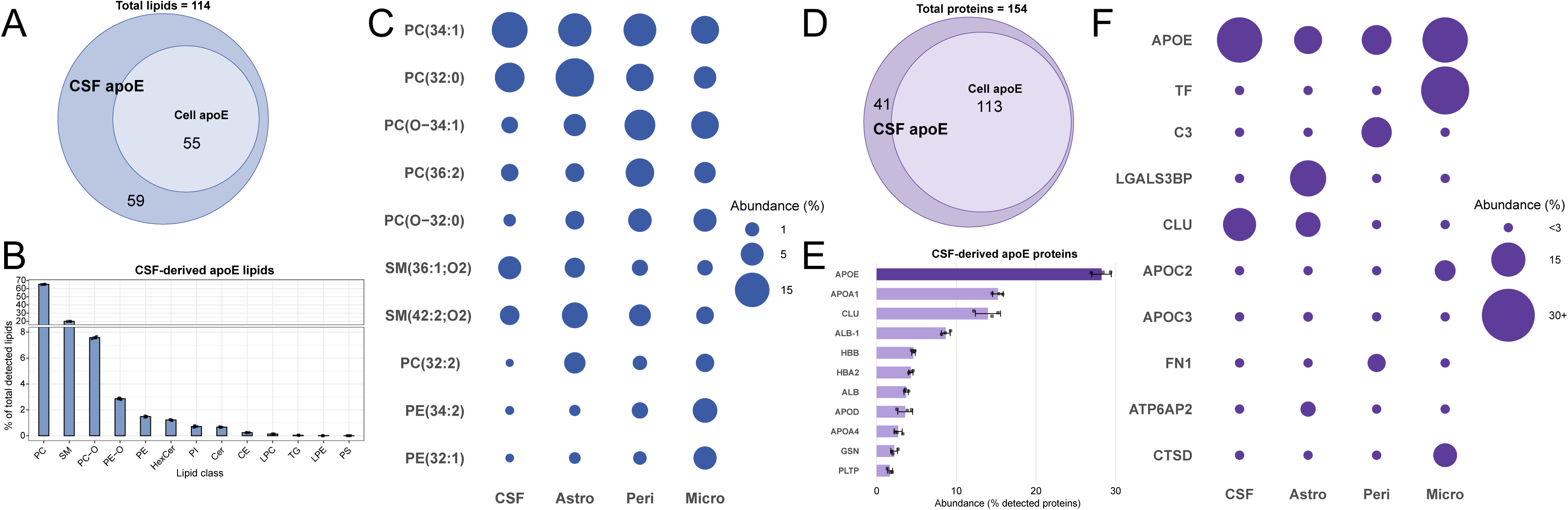
The components of cell-secreted apoE particles’ lipidome and proteotype are found in human CSF. A) The overlap of lipids identified as associated with CSF and cell-secreted apoE. 114 lipids were detected and annotated in the dataset. ‘Cell apoE’ lipids are defined as enriched (log_2_FC ≥ 1, FDR < 0.05) in each condition vs. vehicle media controls. B) Relative abundance of lipid classes in CSF-derived apoE particles. Points in graphs represent individual replicates (n = 3), bars represent the mean, and error bars ± SD. C) Dot plot shows that the most abundant lipid species detected in cell-secreted apoE are also present in CSF apoE. Lipid abundances are expressed as the percentage contribution of each lipid species to the total lipid signal per sample. Mean percentages of three replicates are shown. D) The overlap of proteins identified as associated with CSF and cell-secreted apoE. 154 proteins were detected and validated in the dataset. ‘Cell apoE’ proteins are defined as enriched (log_2_FC ≥ 1, FDR < 0.05) in each condition vs. vehicle media controls. E) Proteins contributing cumulatively to 90% of total CSF apoE proteotype and their relative abundances, expressed as percentage of all detected proteins. Points in graphs represent individual replicates (n = 3), bars represent the mean, and error bars ± SD. F) Dot plot shows that selected (most abundant and biologically relevant) proteins detected in cell-secreted apoE are also present in CSF apoE. Protein abundances are expressed as the percentage contribution of each protein to the total protein signal per sample. The mean percentages of three apoE immunoprecipitation replicates from pooled human CSF are shown.

### 3.5 ApoE particles secreted by brain cells contain lipids and proteins that are found in CSF

After confirming that the composition of brain apoE particles depends on the cell type of origin, we aimed to verify the presence of identified lipids and proteins *in vivo*. To do so, we used apoE particles immunoprecipitated from human CSF as the reference. We identified 114 lipids in CSF apoE, 55 of which were common with cell-secreted apoE particles (Fig. 4 A, Sup. File 2). Relative abundance calculations of lipid classes in CSF-derived apoE particles revealed a high proportion of PC (65.2%), SM (20.0%), and PC-O (7.6%), resembling the typical lipidome of blood-borne HDL particles^32^ and the composition of astrocyte-secreted apoE (Fig. 4 B, Sup. Fig. 3 C, Sup. File 3). We then selected the 10 most prevalent lipids detected in brain cell-secreted apoE particles and compared their relative levels with the lipids of CSF-derived apoE. We observed similar distribution patterns across all conditions but with short-chain PC 32:2 and two PE species being more abundant in cell-than in CSF-derived apoE particles (Fig. 4 C).

We identified 154 proteins associated with CSF apoE, 113 of which were common with cell-secreted apoE particles (Fig. 4 D, Sup. File 1). Calculations of relative abundance showed APOE (28.2 % of total protein signal), other apolipoproteins (APOA1 – 15.2%, CLU – 14.0%, APOD – 3.6%, APOA4 – 2.7%), GSN (2.2%), and PLTP (1.7%) among proteins contributing to 90% of total CSF-derived apoE proteotype. We also noted the presence of albumin (ALB and ALB-1; total 12.4%) and hemoglobin subunits (HBA2 and HBB; total 8.9%) (Fig. 4 E, Sup. File 4). We then selected 10 proteins associated with brain cell-secreted apoE particles, which were among the most abundant and could be linked to relevant biological functions or represented each of the common protein groups detected (lipid metabolism: APOE, APOC2, APOC3, CLU; amyloid processing: CTSD and CLU; extracellular matrix interactions: FN1, LGALS3BP; implication in neurodegenerative diseases – ATP2AP2^33^; complement system component C3; iron transport – TF). We compared their relative levels in brain cell-and CSF-derived apoE particles and observed that these proteins were also present in CSF apoE particles, although their abundances differed (Fig. 4 F). Taken together, these results showed that CSF-derived and brain cell-secreted apoE particles are composed of largely overlapping lipid and protein components, suggesting that despite differences in relative abundance, their molecular compositions are similar.

## 4. Discussion

In this study, we demonstrate that the lipid and protein composition of CNS apoE particles depends on their cellular origin. The protein and lipid profiles of apoE particles secreted by astrocytes, microglia, and pericytes were different, showing that CNS apoE is a diverse population of particles rather than a single, uniform type of lipoprotein. Notably, cell type-specific apoE particles contained proteins and lipids associated with lipid metabolism, oxidative stress resistance, lysosomal function, extracellular matrix biology, iron transport, and amyloid beta processing. This suggests that they may play distinct roles in brain homeostasis and neurodegenerative diseases.

Ether-linked phosphatidylcholines (PC-O) and sphingomyelins (SM) showed the strongest cell type specificity among the differences in particles’ lipid composition. PC-O were enriched in pericyte- and microglia-secreted apoE, while SM predominated in astrocyte-secreted particles. PC-O contribute to membrane rigidity and protect against oxidative damage^34,35^. Along with the enrichment of antioxidant proteins (PRDX1 and PRDX6), the increased presence of PC-O in pericyte-secreted apoE suggests a potential role in oxidative stress resistance. Oxidative stress-related disturbances are implicated in blood-brain barrier (BBB) dysfunction and neurodegeneration^36,37^. Since SM also increase membrane rigidity through interactions with cholesterol and are metabolically co-regulated with PC-O^38^, elevated PC-O may compensate for lower SM levels in pericyte- and microglia-secreted particles. Microglial apoE particles were characterized by higher PE content and lower overall lipidation compared to astrocyte- and pericyte-secreted apoE. PE enrichment, particularly in iPSC-secreted apoE, together with lower phosphatidylcholine abundance, suggests that some apoE may be transported in extracellular vesicles rather than in lipoproteins ^39–42^. These findings underscore the importance of defining the distribution of apoE between extracellular vesicles and lipoproteins across various CNS cell types. Reduced lipidation of microglial apoE can be explained by lower abundance of PLTP, reduced CLU, and increased APOC2, which promotes lipoprotein lipase–mediated triglyceride hydrolysis^43–45^. Low lipidation of apoE particles in microglia observed in our study is consistent with previous reports describing these cells as producers of relatively lipid-poor apoE-containing lipoproteins^46,47^. However, this finding contrasts with more recent evidence demonstrating that murine microglia secrete larger apoE particles than astrocytes^48^. Similarly, a recent study identified an enrichment of cholesteryl esters in microglial apoE^49^. These discrepancies may reflect species-specific differences, as the contrasting observations were obtained from mouse glia, whereas we used human iPSC-derived cells in this study. There are marked differences between human and mouse lipoproteins and microglia, including differences in transcriptional programs, lipid metabolism, and functional responses^50–53^. Among these variations, the regulation of the ATP binding cassette transporter ABCA1 is particularly relevant. ABCA1 is the main transporter responsible for cholesterol and phospholipid transfer to apoE particles, and its regulation differs between human and mouse models^54^. This species-specific regulation of apoE lipidation pathways may explain the differences in apoE particle composition observed between studies. Conversely, higher lipidation of astrocyte- and pericyte-derived apoE may reflect more efficient apoE recycling. This process has been described in peripheral cells and has been proposed to occur in CNS glia^55–58^. The capacity of microglia to recycle apoE and generate more lipidated lipoprotein particles might be of particular interest as recent studies showed that impaired apoE secretion during inflammation or in *APOE4/E4* microglia led to increased intracellular lipid accumulation and microglial dysfunction^59–61^. Alternatively, the observed low lipidation of microglial apoE might be due to the limited sensitivity of untargeted lipidomics methods to cholesterol and cholesteryl esters. Notably, the present study focused on sphingolipid, phospholipid, glycerolipid, and cholesteryl ester classes, which can be reliably profiled together using a single LC-MS/MS method. Given the central role of cholesterol metabolism in neurodegenerative diseases, future studies incorporating cholesterol-optimized methods would complement these findings and provide a more complete picture of apoE particle composition.

We also observed cell type-specific differences in apoE-associated proteins involved in amyloid beta metabolism and lysosomal function. The differential abundance of CLU, CTSD, and ADAM10 suggests that apoE particles from different cellular sources may regulate amyloid processing and clearance differently. Additionally, ATP6AP2, a vacuolar ATPase component required for lysosomal acidification, was enriched in astrocyte-derived apoE, suggesting a potential relationship between astrocyte-derived apoE and lysosomal function^33,62–64^. Our findings are consistent with a recent study by Strickland *et al*., who reported the enrichment of PLTP and CLU in astrocyte-derived apoE and LPL in microglial apoE, as well as increased sphingomyelin content in astrocyte particles^49^. We noted an upregulation of complement system components in pericyte-secreted apoE, consistent with the evidence about complement secretion by pericytes^65^ and the role of the complement cascade in maintaining BBB integrity^66^. CypA, identified as a component of astrocyte- and pericyte-secreted apoE particles, is also involved in the BBB regulation, with increased CypA activity contributing to BBB breakdown^67^. The high contribution of transferrin to microglial apoE suggests a potential role of these particles in iron homeostasis, which has been implicated in inflammation and neurodegenerative diseases^68^.

A few methodological considerations are worth noting. First, a population of incompletely differentiated astrocytes remained after differentiation; however, additional passaging before apoE particle immunoprecipitation and previous reports describing *Sox2* expression in mature astrocytes mitigate concerns about its impact^69^. To address lower apoE secretion by astrocytes, we report relative lipid and protein abundances across investigated cell types. Additionally, in heatmaps and volcano plots comparing proteins associated with apoE-containing lipoproteins, proteomics data were normalized to APOE intensity. Second, our study focused exclusively on *APOE3/E3* cells. While *APOE* genotype is a major determinant of AD risk, recent evidence suggests that apoE lipid composition is more strongly influenced by cell type than by genotype^49^. Lastly, our immunoprecipitation approach could not distinguish between lipidated apoE, lipid-free apoE, and apoE associated with extracellular vesicles.

Including CSF-derived apoE as an *in vivo* reference allowed us to evaluate the physiological relevance of our findings. The lipid composition of CSF-derived apoE resembled that of CNS lipoproteins and was consistent with previous CSF lipidomics studies^70,71^. Similarly, the major proteins associated with CSF-derived apoE overlapped significantly with those previously identified in CSF lipoproteins^14,64,72–77^. Hemoglobin beta reflected minor blood contamination^78^ and apoA-I likely originated from peripheral high-density lipoprotein (HDL) entering the CSF^79–83^, therefore these proteins were excluded from analyses of cell-secreted apoE. Albumin, although potentially reflecting blood contamination, has been previously reported in CSF lipoproteins^14^ and can bind lipids^84^. However, we excluded it from the cell-secreted apoE analyses due to its high presence in vehicle media controls. Most proteins and lipids identified in cell-secreted apoE were also present in CSF-derived apoE, supporting the biological relevance of our *in vitro* model. Features that were only enriched in apoE secreted by cultured cells, such as LGALS3BP and PC 40:5, may represent culture-specific components or transient constituents of newly secreted apoE particles. However, both have previously been implicated in AD pathology^85–87^.

In summary, our findings demonstrate that CNS apoE particles are molecularly heterogeneous and that their protein and lipid composition is determined by their cellular origin. ApoE particles from different cell types differ in molecular features linked to oxidative stress resistance, lipid metabolism, extracellular matrix biology, lysosomal function, and amyloid processing. This suggests that apoE secreted by astrocytes, microglia, and pericytes may perform distinct functions in brain homeostasis and neurodegenerative diseases.

## Supporting information

Sup. File 1. Protein filtering

Sup. File 2. Lipid filtering

Sup. File 3. Lipid relative abundance

Sup. File 4. Protein relative abundance

Sup. Table 1. Lipid membership table

Sup. Table 2. Protein membership table

## 5. Acknowledgements

JR is supported by a grant from the Synapsis Foundation for Dementia Research (No. 2022-PI04) and a grant from the Bright Focus Foundation (A2021037S). Illustrations were made with templates from NIH BioArt Source (https://bioart.niaid.nih.gov/), Bioicons (https://bioicons.com/), and BioRender.com (License Robert_2026). We appreciate the support of the metabolomics team at the Functional Genomics Center Zurich in lipidomics analysis.

## 7. Author contributions

Conceptualization – JR and EV;

Data collection – EV, SG, AJ, SHA, SH;

Data curation and analysis – EV, SG, AJ, AO;

Methodology – EV, SG, AJ, Validation and visualization – EV;

Supervision and funding acquisition – JR and AvE;

Writing of the first draft – EV;

Review and editing – all authors.

## 8. Data availability

The mass spectrometry proteomics data generated during this study have been submitted to the ProteomeXchange Consortium via its partner repository MassIVE, (MassIVE identifier: MSV000102791, ProteomeXchange identifier: PXD082409). The mass spectrometry lipidomics data have been submitted to MassIVE (identifier: MSV000103068).

---

**Supplementary Figure 1.**
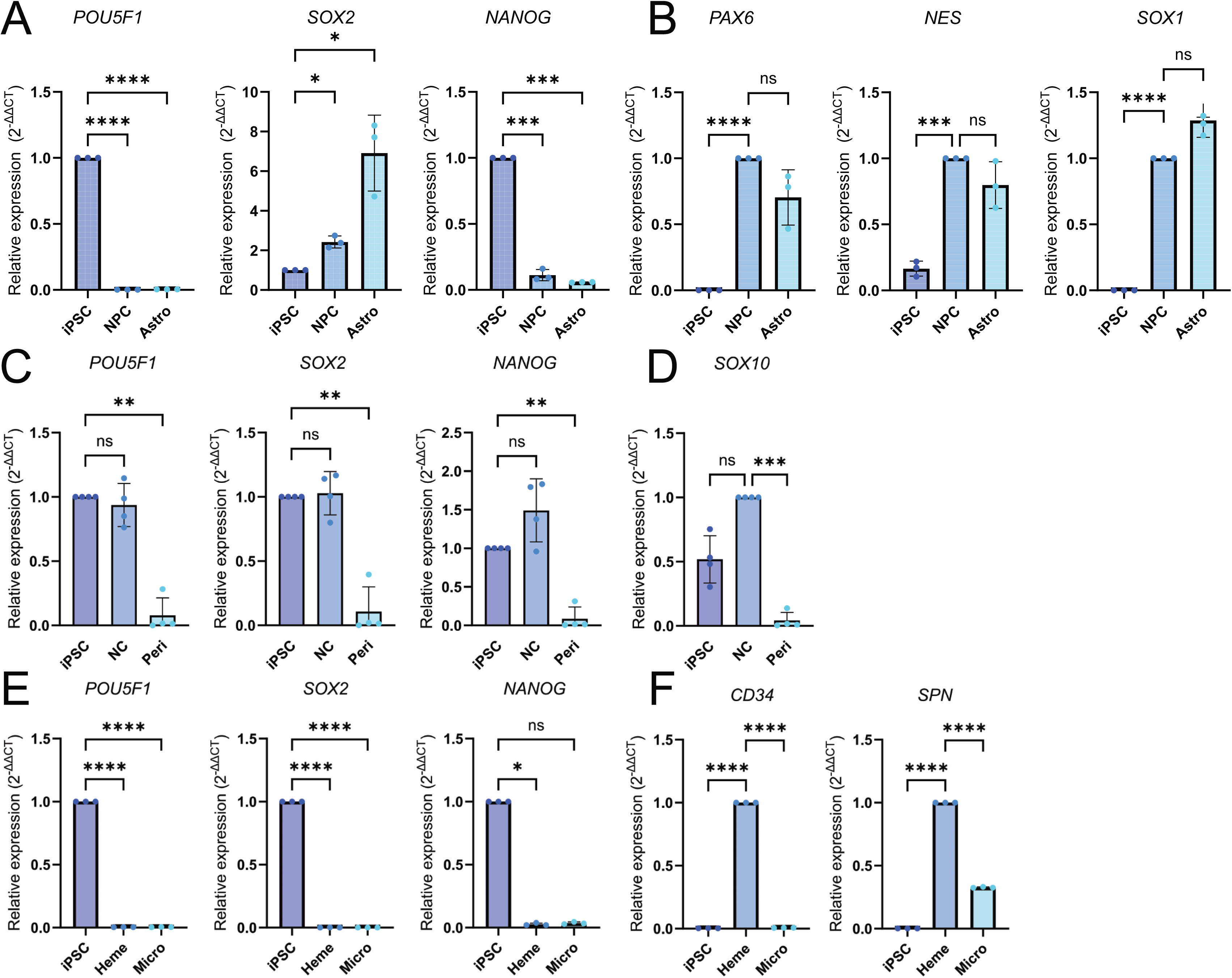
Relative gene expression of pluripotency and intermediate differentiation stage markers across iPSC-derived cell types. A) Gene expression of pluripotency markers (*POU5F1*, *SOX2, NANOG*) and (B) neural progenitor markers (*PAX6, NES, SOX1*) in iPSCs, NPC, and astrocytes. C) Gene expression of pluripotency markers (*POU5F1, SOX2, NANOG*) and (D) the neural crest marker (*SOX10*) in iPSCs, NC, and pericytes. E) Gene expression of pluripotency markers (*POU5F1*, *SOX2, NANOG*) and (F) hematopoietic stem cell markers (*CD34, SPN*) in iPSCs, Heme, and microglia. Points in graphs represent individual experiments (biological replicates, n = 3-4), bars represent the mean, and error bars ± SD, *p < 0.05, **p < 0.01, ***p < 0.001, ****p < 0.0001, ns – not significant. iPSC – induced pluripotent stem cells, NC – neural crest, Heme – hematopoietic stem cells, Peri – pericytes, Micro – microglia, Astro – astrocytes.

**Supplementary Figure 2.**
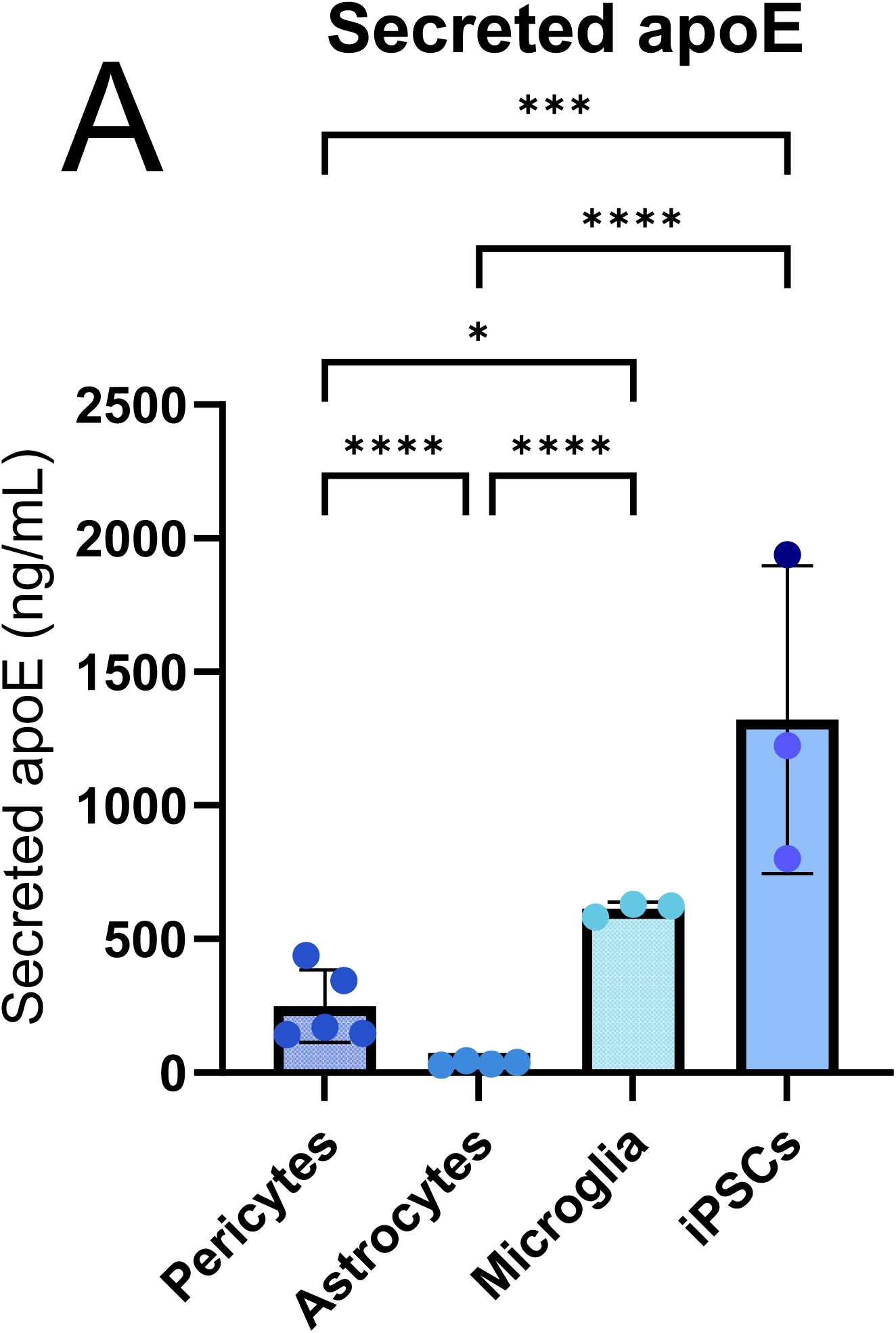
ApoE levels in culture media of iPSC-derived cells. A) ApoE levels in the culture media of each cell type, measured by ELISA. Points in graphs represent individual experiments (biological replicates, n = 3-5), bars represent the mean, and error bars ± SD, *p < 0.05, **p < 0.01, ***p < 0.001, ****p < 0.0001. Only significant pairwise comparisons are shown.

**Supplementary Figure 3.**
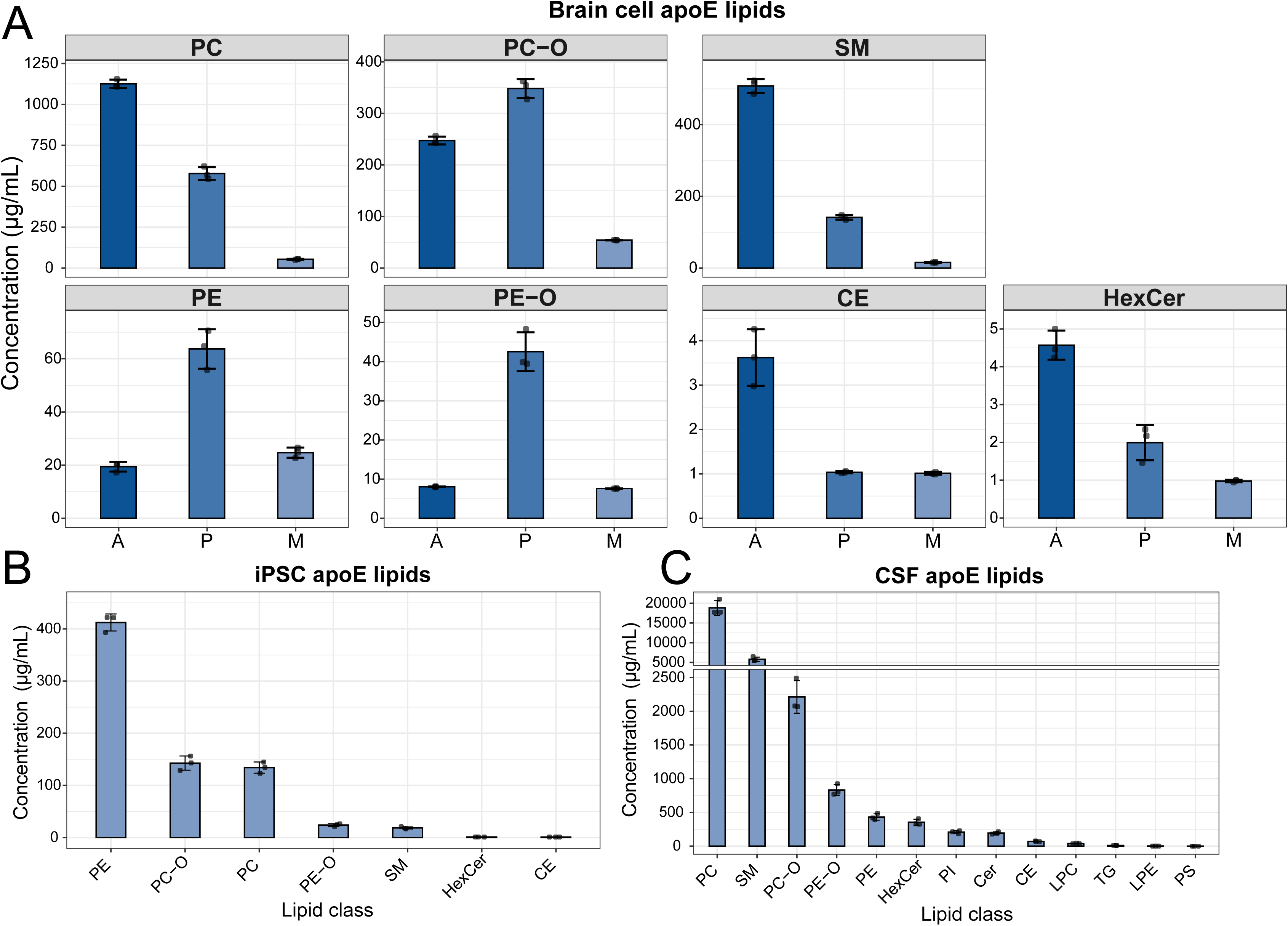
Lipid class concentrations. A) Absolute concentrations of lipid classes in astrocyte-, pericyte-, and microglia-secreted apoE particles. A – astrocyte-, P – pericyte-, M – microglia-secreted apoE. B) Absolute concentrations of lipid classes in iPSC-secreted apoE particles. C) Absolute concentrations of lipid classes in CSF-derived apoE particles. Points in graphs represent individual replicates (n = 3), bars represent the mean, and error bars ± SD.

**Supplementary Figure 4.**
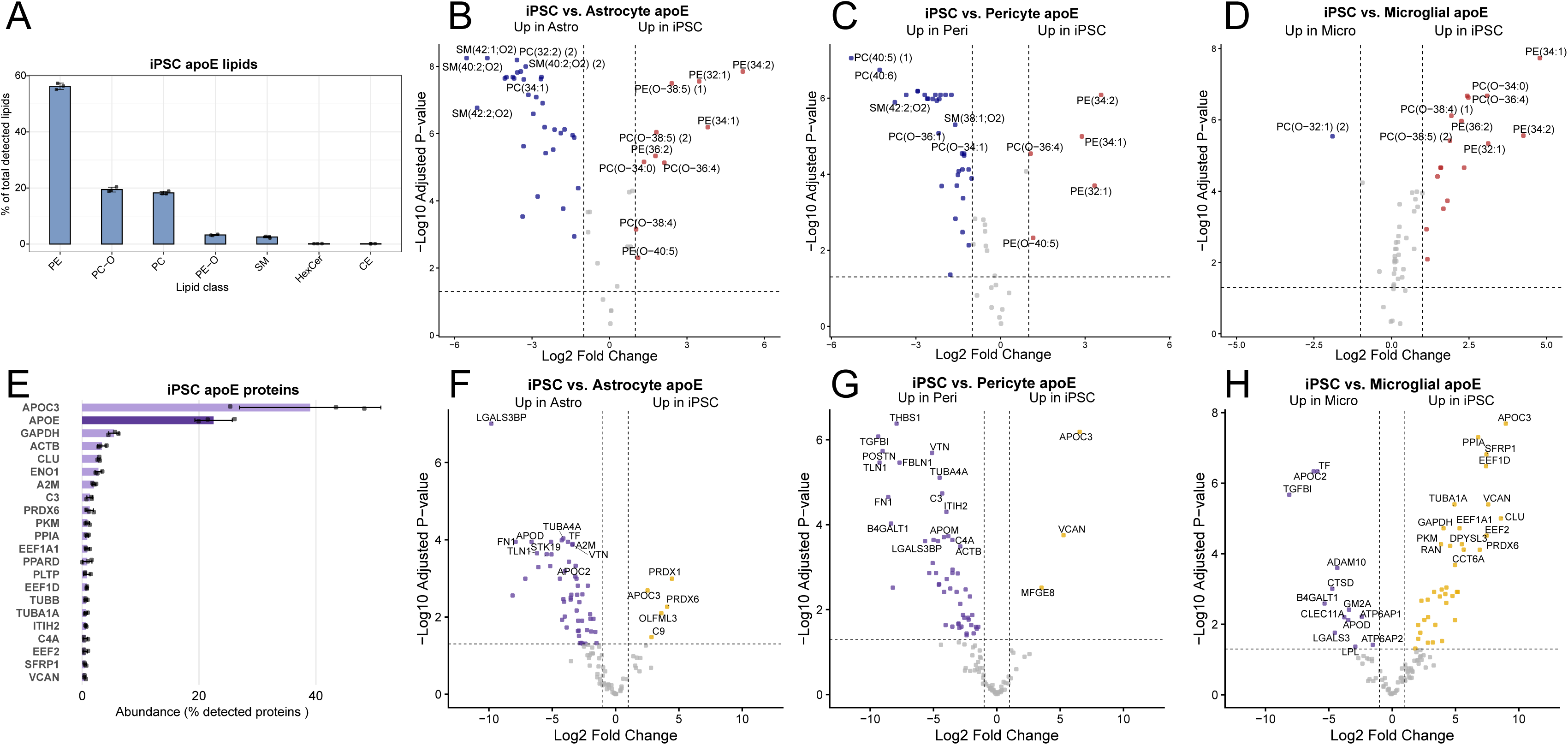
Composition of iPSC-secreted apoE particles. A) Relative abundance of lipid classes in iPSC-secreted apoE particles. Points in graphs represent individual replicates (n = 3), bars represent the mean, and error bars ± SD. B-D) Differential expression of lipids associated with iPSC vs. astrocyte (B), iPSC vs. pericyte (C), and iPSC vs. microglial (D) apoE particles. Each point represents one lipid species. Significance thresholds: |log_2_FC| ≥ 1, FDR < 0.05. E) Proteins contributing cumulatively to 90% of total iPSC-secreted apoE proteotype and their relative abundances, expressed as percentage of all detected proteins. Points in graphs represent individual replicates (n = 3), bars represent the mean, and error bars ± SD. F-H) Differential expression of proteins associated with iPSC vs. astrocyte (F), iPSC vs. pericyte (G), and iPSC vs. microglial (H) apoE particles. Data are normalized to APOE intensity in addition to global (median-centered) normalization. Significance thresholds: |log_2_FC| ≥ 1, FDR < 0.05.

## References

1 Holtzman, D. M., Herz, J. & Bu, G. Apolipoprotein E and apolipoprotein E receptors: normal biology and roles in Alzheimer disease. Cold Spring Harb Perspect Med 2, a006312 (2012). 10.1101/cshperspect.a006312

2 Corder, E. H. et al. Gene dose of apolipoprotein E type 4 allele and the risk of Alzheimer’s disease in late onset families. Science 261, 921–923 (1993). 10.1126/science.8346443

3 Hansen, L. A. et al. Apolipoprotein-E epsilon-4 is associated with increased neurofibrillary pathology in the Lewy body variant of Alzheimer’s disease. Neurosci Lett 182, 63–65 (1994). 10.1016/0304-3940(94)90206-2

4 Scheltens, P. et al. Alzheimer’s disease. Lancet 397, 1577–1590 (2021). 10.1016/S0140-6736(20)32205-4

5 Blumenfeld, J., Yip, O., Kim, M. J. & Huang, Y. Cell type-specific roles of APOE4 in Alzheimer disease. Nature Reviews Neuroscience 25, 91–110 (2024). 10.1038/s41583-023-00776-9

6 Boyles, J. K., Pitas, R. E., Wilson, E., Mahley, R. W. & Taylor, J. M. Apolipoprotein E associated with astrocytic glia of the central nervous system and with nonmyelinating glia of the peripheral nervous system. Journal of Clinical Investigation 76, 1501–1513 (1985). 10.1172/JCI112130

7 Bruinsma, I. B. et al. Apolipoprotein E protects cultured pericytes and astrocytes from D-Aβ1–40-mediated cell death. Brain Research 1315, 169–180 (2010). 10.1016/j.brainres.2009.12.039

8 Keren-Shaul, H. et al. A Unique Microglia Type Associated with Restricting Development of Alzheimer’s Disease. Cell 169, 1276–1290.e1217 (2017). 10.1016/j.cell.2017.05.018

9 Fernandez-Calle, R. et al. APOE in the bullseye of neurodegenerative diseases: impact of the APOE genotype in Alzheimer’s disease pathology and brain diseases. Mol Neurodegener 17, 62 (2022). 10.1186/s13024-022-00566-4

10 Jackson, R. J., Hyman, B. T. & Serrano-Pozo, A. Multifaceted roles of APOE in Alzheimer disease. Nat Rev Neurol 20, 457–474 (2024). 10.1038/s41582-024-00988-2

11 Zheng, J. Y. et al. Selective deletion of apolipoprotein E in astrocytes ameliorates the spatial learning and memory deficits in Alzheimer’s disease (APP/PS1) mice by inhibiting TGF-beta/Smad2/STAT3 signaling. Neurobiol Aging 54, 112–132 (2017). 10.1016/j.neurobiolaging.2017.03.002

12 Henningfield, C. M., Arreola, M. A., Soni, N., Spangenberg, E. E. & Green, K. N. Microglia-specific ApoE knock-out does not alter Alzheimer’s disease plaque pathogenesis or gene expression. Glia 70, 287–302 (2022). 10.1002/glia.24105

13 Yamazaki, Y. et al. ApoE (Apolipoprotein E) in Brain Pericytes Regulates Endothelial Function in an Isoform-Dependent Manner by Modulating Basement Membrane Components. Arterioscler Thromb Vasc Biol 40, 128–144 (2020). 10.1161/ATVBAHA.119.313169

14 Merrill, N. J. et al. Human cerebrospinal fluid contains diverse lipoprotein subspecies enriched in proteins implicated in central nervous system health. Science Advances 9 (2023). 10.1126/sciadv.adi5571

15 Nimsanor, N. et al. Generation of induced pluripotent stem cells derived from a 77-year-old healthy woman as control for age related diseases. Stem Cell Research 17, 550–552 (2016). 10.1016/j.scr.2016.09.019

16 Faal, T. et al. Induction of Mesoderm and Neural Crest-Derived Pericytes from Human Pluripotent Stem Cells to Study Blood-Brain Barrier Interactions. Stem Cell Reports 12, 451–460 (2019). 10.1016/j.stemcr.2019.01.005

17 Fan, J. et al. Small molecule inducers of ABCA1 and apoE that act through indirect activation of the LXR pathway. J Lipid Res 59, 830–842 (2018). 10.1194/jlr.M081851

18 Mellacheruvu, D. et al. The CRAPome: a contaminant repository for affinity purification–mass spectrometry data. Nature Methods 10, 730–736 (2013). 10.1038/nmeth.2557

19 Szklarczyk, D. et al. The STRING database in 2023: protein–protein association networks and functional enrichment analyses for any sequenced genome of interest. Nucleic Acids Research 51, D638–D646 (2023). 10.1093/nar/gkac1000

20 Ganguin, A. A., Skorup, I., Streb, S., Othman, A. & Luciani, P. Formation and Investigation of Cell-Derived Nanovesicles as Potential Therapeutics against Chronic Liver Disease. Advanced Healthcare Materials 12, 2300811 (2023). 10.1002/adhm.202300811

21 McDonald, J. G. et al. Introducing the Lipidomics Minimal Reporting Checklist. Nature Metabolism 4, 1086–1088 (2022). 10.1038/s42255-022-00628-3

22 Nikodemova, M. & Watters, J. J. Efficient isolation of live microglia with preserved phenotypes from adult mouse brain. Journal of Neuroinflammation 9, 635–635 (2012). 10.1186/1742-2094-9-147

23 Goetze, S. et al. Reproducible Determination of High-Density Lipoprotein Proteotypes. Journal of Proteome Research 20, 4974–4984 (2021). 10.1021/acs.jproteome.1c00429

24 Naba, A. et al. The Matrisome: In Silico Definition and In Vivo Characterization by Proteomics of Normal and Tumor Extracellular Matrices. Molecular & Cellular Proteomics 11, M111.014647–M014111.014647 (2012). 10.1074/mcp.M111.014647

25 de Retana, S. F. et al. Peripheral administration of human recombinant ApoJ/clusterin modulates brain beta-amyloid levels in APP23 mice. Alzheimers Res Ther 11, 42 (2019). 10.1186/s13195-019-0498-8

26 Gallwitz, L. et al. Cathepsin D: Analysis of its potential role as an amyloid beta degrading protease. Neurobiology of Disease 175, 105919–105919 (2022). 10.1016/j.nbd.2022.105919

27 Kuhn, P. H. et al. ADAM10 is the physiologically relevant, constitutive α-secretase of the amyloid precursor protein in primary neurons. The EMBO Journal 29, 3020–3032 (2010). 10.1038/emboj.2010.167

28 Zhao, Y. et al. β2-Microglobulin coaggregates with Aβ and contributes to amyloid pathology and cognitive deficits in Alzheimer’s disease model mice. Nature Neuroscience 26, 1170–1184 (2023). 10.1038/s41593-023-01352-1

29 Neniskyte, U. & Brown, G. C. Lactadherin MFG -E8 is essential for microglia-mediated neuronal loss and phagoptosis induced by amyloid β. Journal of Neurochemistry 126, 312–317 (2013). 10.1111/jnc.12288

30 Korvatska, O. et al. Altered splicing of ATP6AP2 causes X-linked parkinsonism with spasticity (XPDS). Human Molecular Genetics 22, 3259–3268 (2013). 10.1093/hmg/ddt180

31 Liao, J. et al. Peroxiredoxin 6 in Stress Orchestration and Disease Interplay. Antioxidants 14, 379–379 (2025). 10.3390/antiox14040379

32 Kontush, A., Lhomme, M. & Chapman, M. J. Unraveling the complexities of the HDL lipidome. J Lipid Res 54, 2950–2963 (2013). 10.1194/jlr.R036095

33 Song, Q., Meng, B., Xu, H. & Mao, Z. The emerging roles of vacuolar-type ATPase-dependent Lysosomal acidification in neurodegenerative diseases. Translational Neurodegeneration 9, 17–17 (2020). 10.1186/s40035-020-00196-0

34 Dean, J. M. & Lodhi, I. J. Structural and functional roles of ether lipids. Protein & Cell 9, 196–206 (2018). 10.1007/s13238-017-0423-5

35 Zoeller, R. A. et al. Plasmalogens as endogenous antioxidants: somatic cell mutants reveal the importance of the vinyl ether. The Biochemical journal 338 **( Pt** **3**), 769–776 (1999).

36 Yemisci, M. et al. Pericyte contraction induced by oxidative-nitrative stress impairs capillary reflow despite successful opening of an occluded cerebral artery. Nat Med 15, 1031–1037 (2009). 10.1038/nm.2022

37 Maccioni, R. Cognitive impairment and Alzheimer’s disease: Links with oxidative stress and cholesterol metabolism. Neuropsychiatric Disease and Treatment **Volume** 4, 715–722 (2008). 10.2147/NDT.S3268

38 Milhas, D., Clarke, C. J. & Hannun, Y. A. Sphingomyelin metabolism at the plasma membrane: implications for bioactive sphingolipids. FEBS Lett 584, 1887–1894 (2010). 10.1016/j.febslet.2009.10.058

39 Cole, L. K., Vance, J. E. & Vance, D. E. Phosphatidylcholine biosynthesis and lipoprotein metabolism. Biochimica et Biophysica Acta (BBA) - Molecular and Cell Biology of Lipids 1821, 754–761 (2012). 10.1016/j.bbalip.2011.09.009

40 Kavianpour, A. A. et al. Phosphatidylethanolamine is a phagocytic ligand implicated in the binding and removal of apoptotic and bacterial extracellular vesicles. Curr Biol 35, 4276–4284 e4275 (2025). 10.1016/j.cub.2025.07.043

41 Ding, M. & Rexrode, K. M. A Review of Lipidomics of Cardiovascular Disease Highlights the Importance of Isolating Lipoproteins. Metabolites 10 (2020). 10.3390/metabo10040163

42 Skotland, T., Hessvik, N. P., Sandvig, K. & Llorente, A. Exosomal lipid composition and the role of ether lipids and phosphoinositides in exosome biology. Journal of Lipid Research 60, 9–18 (2019). 10.1194/jlr.R084343

43 Yazdanyar, A., Yeang, C. & Jiang, X.-C. Role of Phospholipid Transfer Protein in High-Density Lipoprotein– Mediated Reverse Cholesterol Transport. Current Atherosclerosis Reports 13, 242–248 (2011). 10.1007/s11883-011-0172-5

44 Ladu, M. J. et al. Lipoproteins in the Central Nervous System. Annals of the New York Academy of Sciences 903, 167–175 (2000). 10.1111/j.1749-6632.2000.tb06365.x

45 Jong, M. C., Hofker, M. H. & Havekes, L. M. Role of ApoCs in lipoprotein metabolism: functional differences between ApoC1, ApoC2, and ApoC3. Arterioscler Thromb Vasc Biol 19, 472–484 (1999). 10.1161/01.atv.19.3.472

46 Fitz, N. F. et al. Phospholipids of APOE lipoproteins activate microglia in an isoform-specific manner in preclinical models of Alzheimer’s disease. Nat Commun 12, 3416 (2021). 10.1038/s41467-021-23762-0

47 Wolfe, C. M., Fitz, N. F., Nam, K. N., Lefterov, I. & Koldamova, R. The Role of APOE and TREM2 in Alzheimer’s Disease-Current Understanding and Perspectives. Int J Mol Sci 20 (2018). 10.3390/ijms20010081

48 Wang, N. et al. Microglial apolipoprotein E particles contribute to neuronal senescence and synaptotoxicity. iScience 27, 110006 (2024). 10.1016/j.isci.2024.110006

49 Strickland, M. R. et al. Lipidome and proteome of astrocyte and microglia ApoE lipoprotein reveal differences based on cell type and ApoE isoform. Journal of Lipid Research 67, 101000–101000 (2026). 10.1016/j.jlr.2026.101000

50 Zhou, Y. et al. Human and mouse single-nucleus transcriptomics reveal TREM2-dependent and TREM2-independent cellular responses in Alzheimer’s disease. Nat Med 26, 131–142 (2020). 10.1038/s41591-019-0695-9

51 Gordon, S. M. et al. A Comparison of the Mouse and Human Lipoproteome: Suitability of the Mouse Model for Studies of Human Lipoproteins. Journal of Proteome Research 14, 2686–2695 (2015). 10.1021/acs.jproteome.5b00213

52 Huynh, T.-P. V. et al. Lack of hepatic apoE does not influence early Aβ deposition: observations from a new APOE knock-in model. Molecular Neurodegeneration 14, 37–37 (2019). 10.1186/s13024-019-0337-1

53 Maloney, B., Ge, Y. W., Alley, G. M. & Lahiri, D. K. Important differences between human and mouse APOE gene promoters: limitation of mouse APOE model in studying Alzheimer’s disease. Journal of Neurochemistry 103, 1237–1257 (2007). 10.1111/j.1471-4159.2007.04831.x

54 Qiu, Y. et al. Human and mouse ABCA1 comparative sequencing and transgenesis studies revealing novel regulatory sequences. Genomics 73, 66–76 (2001). 10.1006/geno.2000.6467

55 Fan, J. et al. An ABCA1-independent pathway for recycling a poorly lipidated 8.1 nm apolipoprotein E particle from glia. J Lipid Res 52, 1605–1616 (2011). 10.1194/jlr.M014365

56 Fazio, S. The Cell Biology and Physiologic Relevance of ApoE Recycling. Trends in Cardiovascular Medicine 10, 23–30 (2000). 10.1016/S1050-1738(00)00033-5

57 Fazio, S., Linton, M. F., Hasty, A. H. & Swift, L. L. Recycling of Apolipoprotein E in Mouse Liver. Journal of Biological Chemistry 274, 8247–8253 (1999). 10.1074/jbc.274.12.8247

58 Hasty, A. H. et al. The recycling of apolipoprotein E in macrophages. Journal of Lipid Research 46, 1433–1439 (2005). 10.1194/jlr.M400418-JLR200

59 Victor, M. B. et al. Lipid accumulation induced by APOE4 impairs microglial surveillance of neuronal-network activity. Cell Stem Cell 29, 1197–1212.e1198 (2022). 10.1016/j.stem.2022.07.005

60 Haney, M. S. et al. APOE4/4 is linked to damaging lipid droplets in Alzheimer’s disease microglia. Nature 628, 154–161 (2024). 10.1038/s41586-024-07185-7

61 Revanna, J. S. et al. Impaired lipoprotein secretion by APOE4 leads to lysosomal and mitochondrial dysfunction in human microglia. bioRxiv (2026). 10.64898/2026.05.12.724612

62 Avrahami, L. et al. Inhibition of Glycogen Synthase Kinase-3 Ameliorates β-Amyloid Pathology and Restores Lysosomal Acidification and Mammalian Target of Rapamycin Activity in the Alzheimer Disease Mouse Model. Journal of Biological Chemistry 288, 1295–1306 (2013). 10.1074/jbc.M112.409250

63 Şentürk, M., et al. Ubiquilins regulate autophagic flux through mTOR signalling and lysosomal acidification. Nature Cell Biology 21, 384–396 (2019). 10.1038/s41556-019-0281-x

64 Ramirez, A. et al. Hereditary parkinsonism with dementia is caused by mutations in ATP13A2, encoding a lysosomal type 5 P-type ATPase. Nature Genetics 38, 1184–1191 (2006). 10.1038/ng1884

65 Xavier, S. et al. Pericytes and immune cells contribute to complement activation in tubulointerstitial fibrosis. Am J Physiol Renal Physiol 312, F516–F532 (2017). 10.1152/ajprenal.00604.2016

66 Alexander, J. J. Blood-brain barrier (BBB) and the complement landscape. Mol Immunol 102, 26–31 (2018). 10.1016/j.molimm.2018.06.267

67 Bell, R. D. et al. Apolipoprotein E controls cerebrovascular integrity via cyclophilin A. Nature 485, 512–516 (2012). 10.1038/nature11087

68 Urrutia, P. J., Borquez, D. A. & Nunez, M. T. Inflaming the Brain with Iron. Antioxidants (Basel) 10 (2021). 10.3390/antiox10010061

69 Wang, Y. et al. SOX2 is essential for astrocyte maturation and its deletion leads to hyperactive behavior in mice. Cell Reports 41, 111842–111842 (2022). 10.1016/j.celrep.2022.111842

70 Byeon, S. K. et al. Cerebrospinal fluid lipidomics for biomarkers of Alzheimer’s disease. Mol Omics 17, 454–463 (2021). 10.1039/d0mo00186d

71 Saito, K. et al. Profiling of Cerebrospinal Fluid Lipids and Their Relationship with Plasma Lipids in Healthy Humans. Metabolites 11 (2021). 10.3390/metabo11050268

72 Borghini, I., Barja, F., Pometta, D. & James, R. W. Characterization of subpopulations of lipoprotein particles isolated from human cerebrospinal fluid. Biochimica et Biophysica Acta (BBA) - Lipids and Lipid Metabolism 1255, 192–200 (1995). 10.1016/0005-2760(94)00232-N

73 Koudinov, A. R., Berezov, T. T., Kumar, A. & Koudinova, N. V. Alzheimer’s amyloid beta interaction with normal human plasma high density lipoprotein: association with apolipoprotein and lipids. Clin Chim Acta 270, 75–84 (1998). 10.1016/s0009-8981(97)00207-6

74 Demeester, N. et al. Characterization and functional studies of lipoproteins, lipid transfer proteins, and lecithin:cholesterol acyltransferase in CSF of normal individuals and patients with Alzheimer’s disease. Journal of Lipid Research 41, 963–974 (2000). 10.1016/S0022-2275(20)32039-3

75 Koch, S. et al. Characterization of four lipoprotein classes in human cerebrospinal fluid. Journal of Lipid Research 42, 1143–1151 (2001). 10.1016/S0022-2275(20)31605-9

76 Suzuki, T. et al. Predominant apolipoprotein J exists as lipid-poor mixtures in cerebrospinal fluid. Annals of clinical and laboratory science 32, 369–376 (2002).

77 Kay, A. D. et al. Remodeling of Cerebrospinal Fluid Lipoprotein Particles after Human Traumatic Brain Injury. Journal of Neurotrauma 20, 717–723 (2003). 10.1089/089771503767869953

78 Barkovits, K. et al. Blood Contamination in CSF and Its Impact on Quantitative Analysis of Alpha-Synuclein. Cells 9, 370–370 (2020). 10.3390/cells9020370

79 Lee, C. Y. D., Tse, W., Smith, J. D. & Landreth, G. E. Apolipoprotein E Promotes β-Amyloid Trafficking and Degradation by Modulating Microglial Cholesterol Levels. Journal of Biological Chemistry 287, 2032–2044 (2012). 10.1074/jbc.M111.295451

80 Timmins, J. M. et al. Targeted inactivation of hepatic Abca1 causes profound hypoalphalipoproteinemia and kidney hypercatabolism of apoA-I. Journal of Clinical Investigation 115, 1333–1342 (2005). 10.1172/JCI23915

81 Elliott, D. A., Weickert, C. S. & Garner, B. Apolipoproteins in the brain: implications for neurological and psychiatric disorders. Clinical Lipidology 5, 555–573 (2010). 10.2217/clp.10.37

82 Stukas, S. et al. Intravenously Injected Human Apolipoprotein A-I Rapidly Enters the Central Nervous System via the Choroid Plexus. Journal of the American Heart Association 3 (2014). 10.1161/jaha.114.001156

83 Kakava, S. et al. Both the low-density lipoprotein receptor and apolipoprotein E define blood-borne high-density lipoprotein entry into the brain. Bioxrive (2025). 10.1101/2025.05.23.655828

84 van der Vusse, G. J. Albumin as fatty acid transporter. Drug Metab Pharmacokinet 24, 300–307 (2009). 10.2133/dmpk.24.300

85 Boza-Serrano, A. et al. Galectin-3 is elevated in CSF and is associated with Abeta deposits and tau aggregates in brain tissue in Alzheimer’s disease. Acta Neuropathol 144, 843–859 (2022). 10.1007/s00401-022-02469-6

86 Yip, P. K. et al. Elevated cerebrospinal fluid galectin-3 and associated cytokines after severe traumatic brain injury in patients. Medicine (Baltimore) 103, e38620 (2024). 10.1097/MD.0000000000038620

87 Castroflorio, E. et al. Lipidomic Analysis of Human Plasma and Hippocampus Across Alzheimer’s Progression and Preclinical 5xFAD Mouse Model. Mol Neurobiol 63 (2026). 10.1007/s12035-026-05849-1

